# 3D *in vitro* blood-brain-barrier model for investigating barrier insults

**DOI:** 10.1101/2022.09.12.507522

**Authors:** Wei Wei, Fernando Cardes, Andreas Hierlemann, Mario M. Modena

## Abstract

The blood-brain-barrier (BBB) prevents that harmful substances in the blood enter the brain, and barrier disruption has been associated with a variety of central-nervous-system diseases. *In vitro* BBB models enable to recapitulate the BBB behavior in a controlled environment to investigate how the barrier reacts to stress events and external insults. Here, we present a human-cell-based BBB platform with integrated, transparent electrodes to monitor barrier tightness in real time at high spatiotemporal resolution. The BBB model includes human cerebral endothelial cells and primary human pericytes and astrocytes in a three-dimensional arrangement within a pump-free, open microfluidic platform. With our platform, we demonstrate that oxygen-glucose deprivation (OGD), which mimics the characteristics of an ischemic insult, induces a rapid remodeling of the cellular actin structures and subsequent morphological changes in the endothelial cells. High-resolution live imaging showed the formation of large actin stress-fiber bundles in the endothelial layer during OGD application, which ultimately led to cell shrinkage and barrier breakage. Simultaneous electrical measurements showed a rapid decrease of the barrier electrical resistance before the appearance of the stress fibers, which indicates that the barrier function is compromised already before the appearance of drastic morphological changes. The results demonstrate that our BBB platform recapitulates the main barrier functions *in vitro* and can be used to investigate the rapid reorganization of the BBB upon application of external stimuli.

## Introduction

The blood-brain barrier (BBB) is a specialized system of brain microvasculature that tightly regulates the transfer of nutrients, metabolites and ions between the blood circulation system and the central nervous system (CNS) to maintain brain homeostasis.[1–4] The high selectivity of the barrier protects the brain from potentially toxic substances and pathogens in the blood, and enables the removal of waste products from the brain, thereby acting as the first-line protection of the CNS. Insults to the BBB, such as strokes and traumatic brain injuries, drastically affect the function and tightness of the BBB, which, in turn, has direct effects on the CNS.[5–8] Furthermore, the high selectivity of the BBB also represents a major roadblock in the effective delivery of drugs and therapeutics to treat CNS diseases,[9–11] including neurodegenerative diseases, cerebrovascular diseases, and brain tumors.[4, 12]

To improve the understanding of the BBB transport mechanisms and its response to external stress events, various static and dynamic *in vitro* BBB models have been reported in literature.[13–22] These models aim at recapitulating morphological and biochemical characteristics of the BBB under controllable and reproducible conditions.[2, 23] Compared to using *in vivo* animal models, *in vitro* models offer lower costs, higher throughput, and the possibility to use human-based cellular models, which reduces potential issues related to specie-dependent differences between human and animal BBBs. Additionally, *in vitro* models have been instrumental in reducing animal use to address ethical concerns.[24–26]

The most commonly used *in vitro* BBB models rely on transwell systems, which include the static culturing of an endothelial-cell barrier on a porous membrane that is immersed in a medium within a well.[13, 14, 16] The simple parallelization of the culture system and its compatibility with a well-plate format have greatly promoted the use of this approach. However, static culturing in a well plate cannot recapitulate important morphological and physiological aspects of the BBB environment, such as liquid flow and its induced shear stress on the endothelial cells.[17, 27] Static cultures may result in BBB models with lower expression of junction proteins, which ultimately affect the barrier structure and function, which, in turn, may lead to poor correlation with *in vivo* data.[23, 28–31] To improve the physiological relevance of *in vitro* models, dynamic *in vitro* BBBs featuring medium flow on the cell monolayer have been developed.[17, 18, 20, 27] Medium perfusion enables improved delivery of oxygen and nutrients to the cells and provides physiological shear stress to the cells,[32] which has been shown to greatly improve the tightness and function of BBB models in comparison to static culturing.[23, 33] Furthermore, microfluidic-based, 3-dimensional (3D) BBB models, which include multiple cell types that are cocultured in a 3D arrangement and often within a hydrogel matrix to better mimic the cellular microenvironment, have been reported in literature.[17, 18, 21, 34] These systems are aimed at overcoming the limitations and low physiological relevance of culturing cells in planar arrangements by providing a 3D microenvironment, where different cell types can communicate, interact and exchange molecular cues with each other. Electrodes were also integrated into microfluidic devices to continually measure the transendothelial electrical resistance (TEER) of the barrier layer as a mean to reliably assess the formation, integrity, and permeability of the endothelial barrier without causing cellular and barrier damage.[35–38] For reliable TEER evaluation, the uniformity of the current density across the cell layer is of fundamental importance.[35, 37] This condition cannot be met by conducting TEER measurements using insertable, thin, chopstick-like electrodes, which feature only low uniformity in current density. Moreover, electrode position may differ between measurements, which further compromises the reliability of the obtained TEER values.[35] Microfluidic devices, featuring integrated TEER sensors with improved current-density profiles, would provide a solution to overcome these limitations.[35, 36]

In this study, we developed an open-microfluidic 3D BBB model and platform, which includes eight BBB devices per plate to perform parallel measurements under various conditions. Each BBB device consisted of a “brain” unit containing primary human astrocytes and human pericytes in a 3D hydrogel, to mimic the brain extracellular matrix (ECM), and a “vascular” microfluidic channel, which was lined with human cerebral endothelial cells. The on-chip vascular unit was exposed to a gradually increasing, quasi-unidirectional flow that was obtained by asymmetric periodic tilting of the BBB platform. The tilting-induced flow promoted the formation of a robust and tight endothelial barrier layer. An integrated indium tin oxide (ITO) - Platinum (Pt), 4-electrode TEER sensor enabled real-time TEER and optical monitoring of the BBB. TEER measurements were used to monitor the formation and dynamic changes of the barrier, while the highly transparent ITO electrodes enabled unrestricted optical access to the barrier model for high-resolution live imaging.

To evaluate the performance of the BBB system and to test for its ability to recapitulate barrier responses to traumatic events, we applied oxygen/glucose deprivation (OGD) conditions to mimic cerebral ischemia. We then investigated the rapid transformations that the BBB undergoes as a result of such an insult. A key pathophysiological feature of cerebral ischemia is a rapid disruption of the BBB, however, its mechanism still remains elusive.[5, 39] The limited understanding of ischemia-induced BBB breakage impedes the development of therapeutic treatments for BBB stabilization, which are aimed at reducing brain edema and neurological damage.[39–41] Our results show that the developed BBB-on-chip platform enables to recapitulate *in vitro* the rapid reorganization of the BBB and endothelial cell cytoskeleton and to monitor it continuously at high spatiotemporal resolution.

## Results

### Device design and fabrication

The blood-brain barrier includes multiple cell types that jointly form a tight and selective barrier structure (Fig. 1a). To reconstitute the BBB structure on chip, we co-cultured human cerebral microvascular endothelial cells (ECs), human astrocytes (HAs) and human pericytes (HPs) in a multi-layer, open microfluidic device (Fig. 1b). ECs were seeded along the walls of a microfluidic channel to form a monolayer structure and recapitulate a microvessel wall. The microchannel was flanked by two open reservoirs to generate gravity-driven flow by tilting the platform, and it was separated from the brain compartment, where HAs and HPs were cultured, by a PET porous membrane. HAs and HPs in the brain compartment were embedded in a hydrogel matrix to provide physical support to the cells and recapitulate the 3-dimensional cellular arrangement. Nutrients in the brain compartment were delivered by transport and diffusion across the endothelial barrier structure and through the opening of the hydrogel/medium reservoir. The open-microfluidic device enabled simple access for sampling on both sides of the cellular barrier to measure molecular transport and diffusion across the BBB. Integrated ITO and Pt electrodes on both sides of the barrier enabled the measurement of TEER values for real-time monitoring of barrier formation and integrity. The transparent, integrated ITO electrodes were patterned on the bottom glass coverslip to enable live, high-resolution confocal microscopy of the BBB structure and its reorganization (Fig. 1c).

**Fig. 1.**
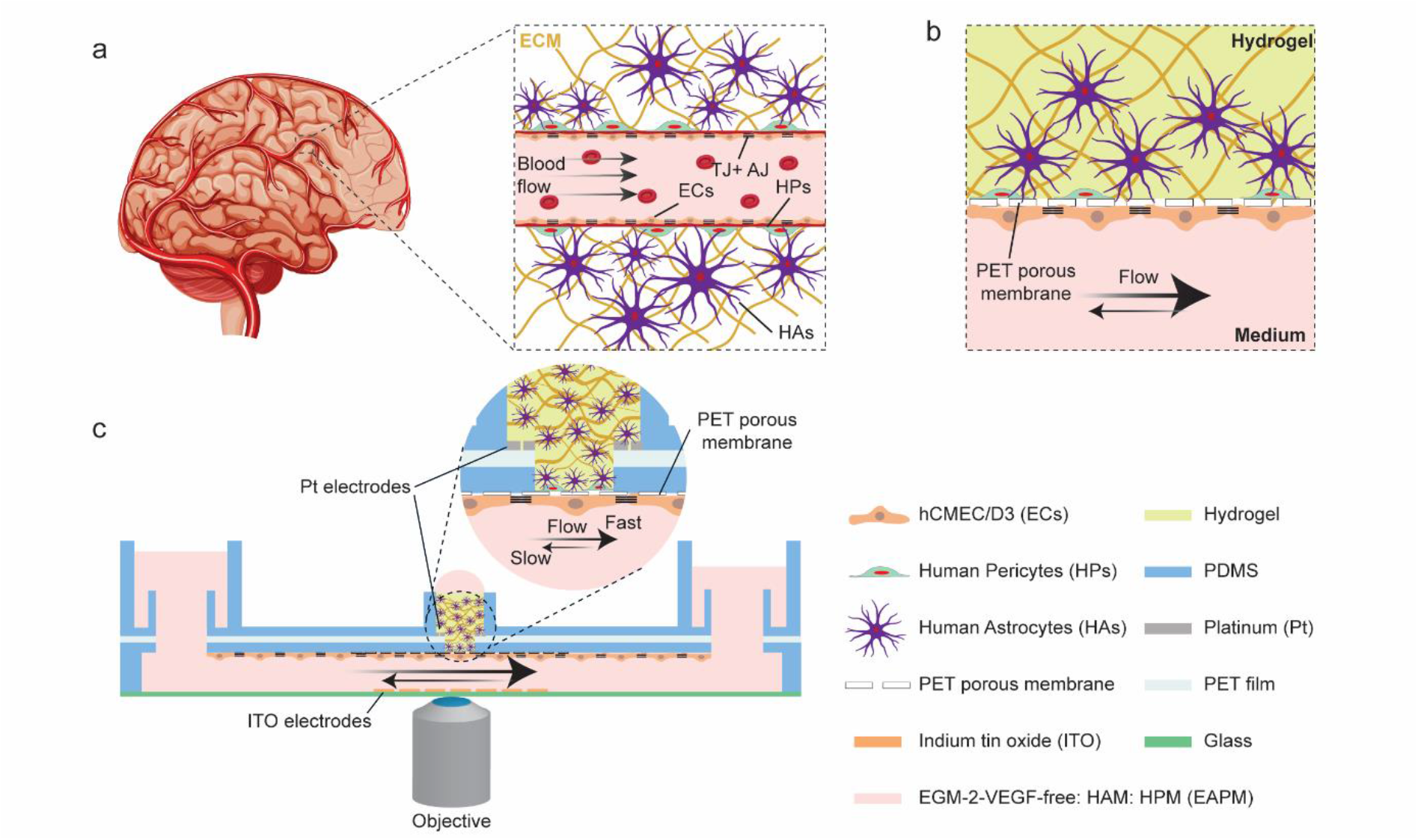
a) Schematic representation of the *in vivo* BBB, in which brain endothelial cells (ECs) line the blood vessels and are exposed to continuous blood flow. Human pericytes (HPs) and astrocytes (HAs) in the extracellular matrix (ECM) surrounded the vessels and, together with the ECs, formed the BBB. Tight junctions (TJ) between endothelial cells limited the transport of molecules across the barrier; b) Schematic representation of the BBB on chip. ECs were seeded on a porous membrane to form a continuous, tight monolayer. Medium flow in the microfluidic channel exposed the ECs to shear stress. HAs and HPs were cultured in a 3D hydrogel structure on the opposite side of the porous membrane; c) Illustration of the BBB chip and its operation. The ECs were cultured in the vascular microchannel, while HAs and HPs were cultured in the upper brain compartment. The coculture medium (EAPM, VEGF-free EGM-2: HAM: HPM = 1: 1: 1) was perfused through the microchannel by gravity-driven flow to deliver oxygen and nutrients to the cells and to expose the EC layer to shear stress. High-resolution microscopy could be carried out through the integrated ITO electrodes, while TEER values could be recorded using the on-chip electrodes to measure barrier tightness.

Each BBB model was realized on a 26 mm × 25.5 mm × 4.2 mm (*L × W × H*) microfluidic device, and the overall BBB platform contained up to 8 devices operated in parallel (Fig. 2a). The microfluidic network included a 100-μm-high microchannel with a central 3.5 mm-diameter BBB region, which was connected to two open reservoirs at each end of the microchannel to realize the vascular unit, and a 3 mm-diameter compartment to host the hydrogel matrix to host the brain compartment (Fig. 2a, b and Supplementary Fig. 1). A porous membrane separated the brain and the vascular side. Tilting of the chip generated a pulsed flow in the vascular channel to expose the ECs lining the channel walls and the porous membrane to shear stress. An asymmetrical tilting scheme was used to generate larger shear stress in one flow direction to avoid symmetric, bidirectional flow that could affect the endothelial layer.[17, 42] The central BBB region on the vascular side was designed with a larger diameter with respect to the top brain compartment so as to expose the ECs forming the BBB to uniform shear stress and to avoid potential barrier leakage caused by lower shear stress along the channel edges (Supplementary Fig. 2). Each system comprised a multi-layer structure of glass-ITO, a thin PDMS layer for microchannel fabrication, a porous PET membrane, an interconnecting PDMS layer to form the brain chamber, a Pt-patterned PET foil, and a final PDMS layer featuring the medium reservoirs and circular rims for stable hanging-drop operation during cell seeding (Fig. 2b). Large Pt contact pads were patterned on the glass-ITO electrodes and on the top PET foil to provide reliable electrical connections and low overall electric resistance. Electrical connections between the printed circuit board (PCB) and the microfluidic devices were realized by spring-loaded pins that contacted the electrode pads from above (Supplementary Fig. 3). Up to 8 chips could be mounted between a custom-made PCB and the chip holder. This arrangement enabled to electrically interrogate the integrated TEER electrodes and holding the chips in position during culturing and imaging. The PCB and the chip holder featured standard well-plate dimensions for compatibility with laboratory automation and microscopy tools. Open windows above and below the microfluidic networks provided optical access for widefield and confocal microscopy (Fig. 2a, Supplementary Fig. 4 and Supplementary Fig. 5). The pump-and tubing-free operation of the chips promoted scalability and parallelization by simple stacking of multiple plates on a tilting stage.

**Fig. 2.**
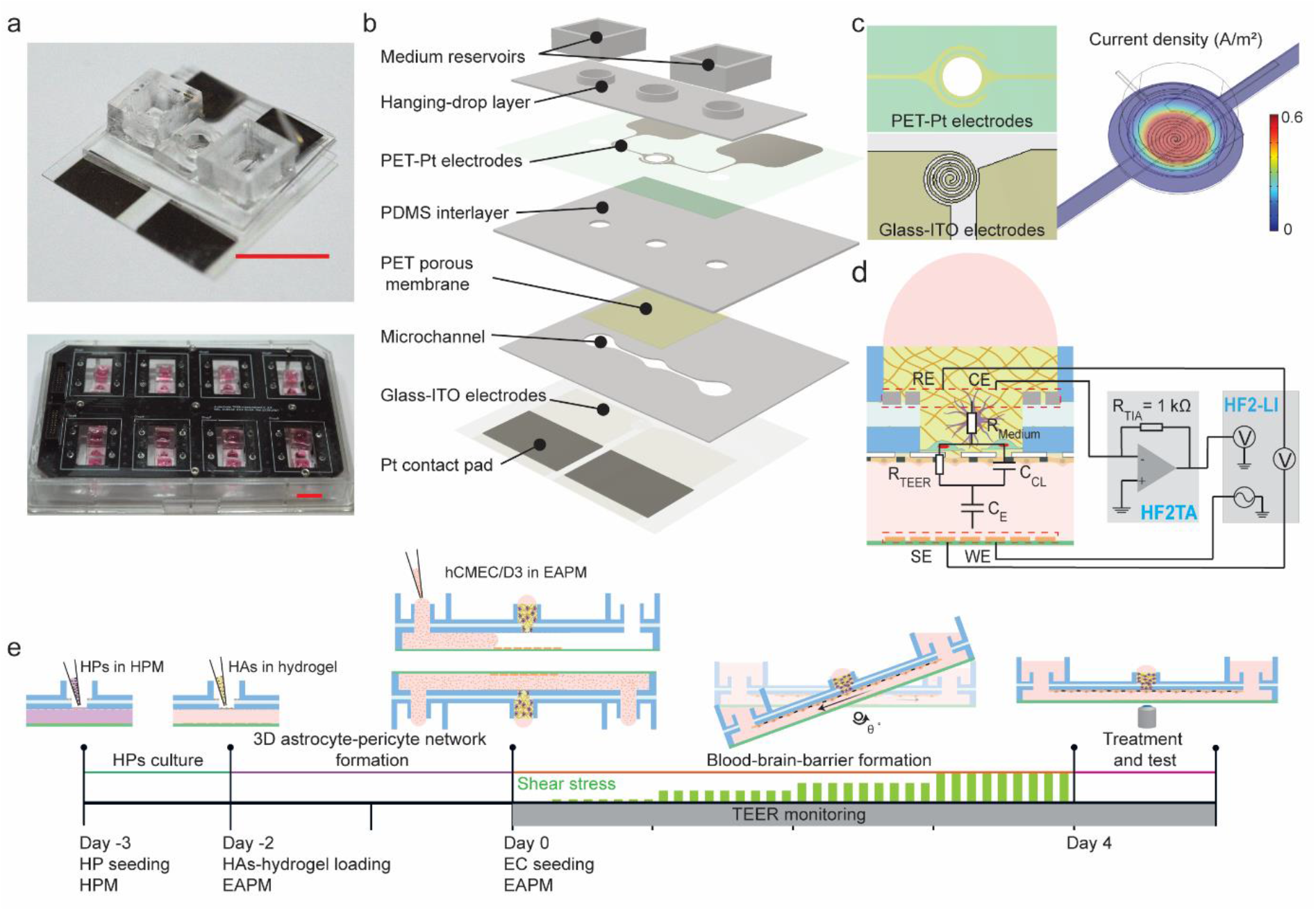
a) Photograph of the BBB chip (top) and the 8-chip BBB platform (bottom). Scale bars = 1 cm; b) Exploded view of the BBB chip, which consists of a glass-ITO coverslip with patterned electrodes, a vascular microchannel layer, a PET porous membrane, a PDMS interlayer, a PET-Pt electrode layer, the hanging-drop layer and two open reservoirs; c) Design of the on-chip, integrated electrodes and FEM simulation of the current density at the EC layer. The glass-ITO electrode was designed in a spiral shape to provide uniform current density at the BBB layer when combined with the top Pt ring electrode. The current density distribution was estimated by simulating a sinusoidal voltage stimulation of 200 mV_p_ amplitude at 1 kHz using COMSOL Multiphysics. The figure shows the current density at the EC layer; d) Schematic design of the 4-electrode TEER measurement system and the electric-equivalent circuit of the BBB (WE: working electrode, SE: sensing electrode, RE: reference electrode, CE: counter electrode), which includes the capacitance of the cell membranes (C_CL_), the resistance of the paracellular route (R_TEER_), the medium and hydrogel resistance (R_medium_), and the electrode capacitances (C_E_); e) Experimental timeline of cell seeding and culturing on the BBB chips. On day -3, HPs were seeded on the porous membrane in the brain compartment, and the device was filled with pericyte medium (HPM). On day -2, HAs, which were previously resuspended in liquid hydrogel solution, were loaded into the brain compartment, and the medium in the device was replaced with the common medium EAPM. On day 0, the ECs were seeded into the microchannel. The platform was then turned upside down and operated in a hanging-drop configuration to promote the sedimentation and the adhesion of the ECs onto the porous membrane. After 2 hours, the platform was returned to a standing-drop configuration, and asymmetric tilting was started to recirculate the medium and apply shear stress to the cell layer. The tilting angle and the resulting shear stress were gradually increased over the 4 days of culturing to obtain a tight cellular barrier. After the barrier had formed, different analyses and treatments were used to assess the integrity and functionality of the BBB model. TEER values were monitored and recorded during the whole EC-barrier formation process.

To obtain a highly uniform current density at the barrier, a spiral-shape bottom electrode was designed to compensate for the low current-density uniformity arising from the top ring-electrodes (Fig. 2c and Supplementary Fig. 3). We employed a four-electrode measurement scheme to eliminate the resistance contributions of the connecting wires and traces and to deal with the high impedance of the double-layer capacitance at the electrode-electrolyte interface. We performed frequency-sweep measurements using a lock-in amplifier to measure the current flowing through the system (voltage-to-current conversion by means of a transimpedance amplifier (TIA)) and the differential voltage across the barrier model (Fig. 2d and Supplementary Fig. 5).

The on-chip BBB was realized by loading the different cell types over multiple days (Fig. 2e). First, HPs were loaded into the brain compartment and left to attach and grow on the porous membrane during 24 h. The following day, the HA-hydrogel mixture was loaded into the brain compartment. The 3D astrocyte-pericyte network formed within 2 days, then, the ECs were finally seeded into the microchannel. To promote adhesion of the ECs to the porous membrane, the chips were maintained in an inverted (hanging-drop) configuration for 2 hours. 3.5-mm-diameter circular hydrophobic rims were realized inside the medium reservoirs to generate a stable hanging-drop network for inverted operation and to prevent medium spillage. After cell attachment, the chips were turned back into their standing-drop configuration, and additional medium was added to the reservoirs to break the liquid drops and to wet the whole reservoir. This procedure was applied to avoid high Laplace pressures of small liquid drops, which would counteract the gravity-induced pressure differences during tilting and reduce the flow rate during barrier formation. The culture medium was continuously recirculated between the two reservoirs of each chip by tilting the platform perpendicular to the microchannel axis. (Supplementary Fig. 6), which provided shear stress to the ECs and increased the gas and nutrient exchange of the medium. The maximum shear stress, exerted on the ECs layer, was gradually increased over the culturing period by increasing the maximum tilting angle to +30° during the 4 days of EC culturing. The platform motion and tilting intervals were optimized to reduce medium backflow so as to generate directional flow across the ECs within physiological shear stress levels[32, 43] (Supplementary Fig. 6).

### BBB formation and characterization

To investigate the efficiency of our system in supporting the growth of different cell types simultaneously, we measured the viability of ECs, Has, and HPs cells using either tissue-specific medium formulation, namely VEGF-free EGM-2, HAM, HPM, or a mixture of them, which we termed EAPM (VEGF-free EGM-2: HAM: HPM = 1: 1: 1). We confirmed that the mixed co-culture medium formulation, EAPM, did not affect the viability of the different cell types in culture. After 2 days of culturing in EAPM, the cell metabolic activity of the HAs (97.12 ± 2.48%) and HPs (94.74 ± 1.93%) did not show any significant difference, in comparison to the viability in their specific medium formulations (Fig. 3a). In contrast, the viability of the ECs was significantly promoted in EAPM (110.13 ± 1.42%) in comparison to the vascular endothelial growth factor (VEGF)-free EGM-2 (Fig. 3b). The high viability of all cell types in the common medium formulation indicated that the cells could indeed be cocultured to reproduce a human BBB model on chip.

**Fig. 3.**
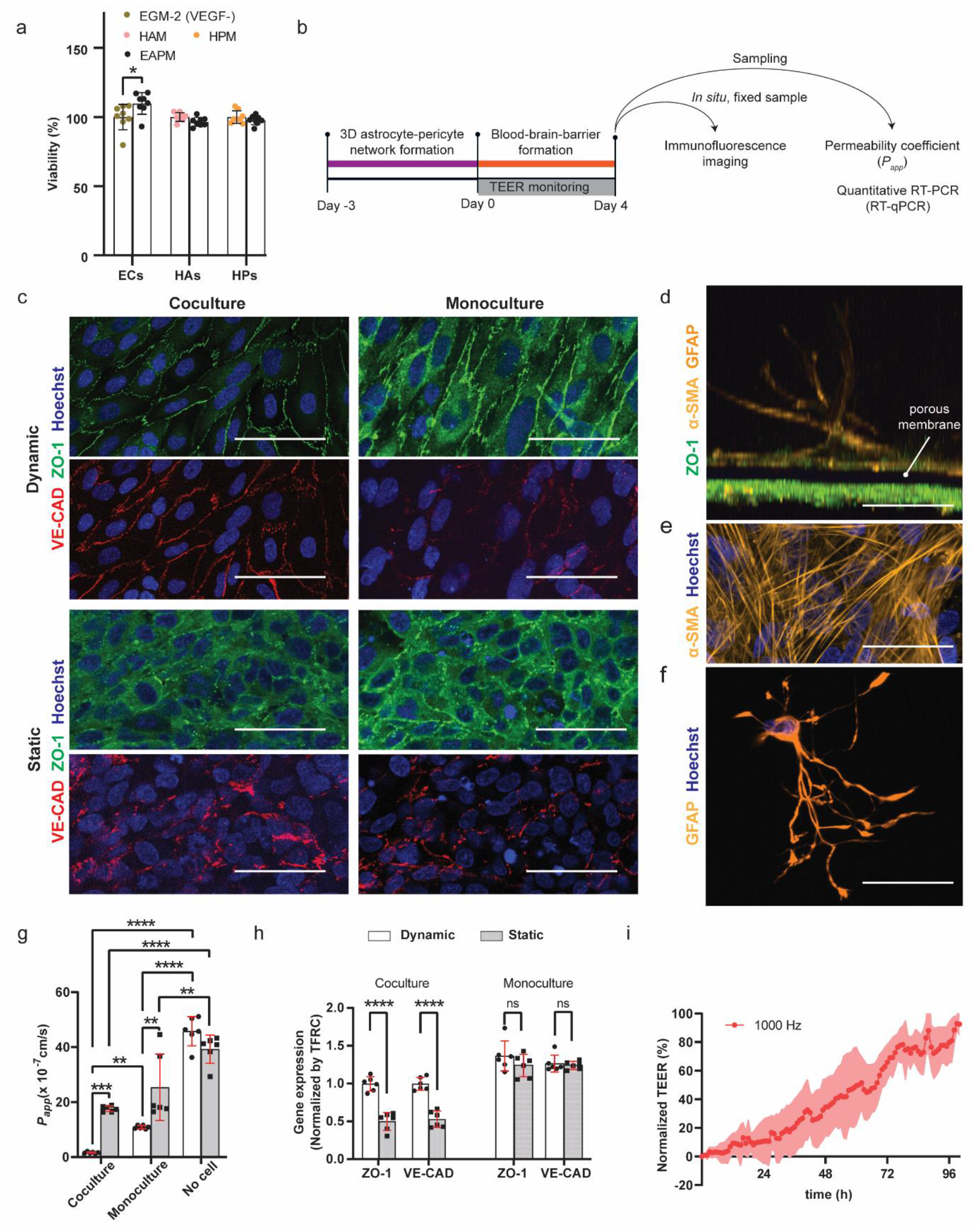
a) Cell viability in specific and common medium formulations (EGM-2 (VEGF-): VEGF-free endothelial medium; HAM: human astrocyte medium; HPM: human pericyte medium; EAPM corresponds to EGM-2-(VEGF-): HAM: HPM = 1: 1: 1; n = 6 for each condition); b) Schematic of the experimental workflow for the characterization of the BBB models; c) Immunofluorescence microscopy images of the EC monolayer under different culturing conditions. The monolayer was stained for the tight-junction protein ZO-1 (green), adherent-junction protein VE-CAD (red) and cellular nuclei (Hoechst, blue). For the dynamic culturing conditions, flow was directed along the vertical direction. However, in the barrier region, flow direction presents different direction components because of the circular compartment and the widening of the channel; d) Vertical cross-section of the BBB model. Pericytes were cultured on the opposite side (top side) of the porous membrane with respect to the EC layer (below the porous membrane). The HPs were stained for α-SMA (yellow) and nuclei (Hoechst, blue). Astrocytes were suspended in the 3D hydrogel matrix and stained for GFAP (yellow). e) Pericytes on the porous membrane; f) Maximum-intensity-projection of an astrocyte on chip in the hydrogel matrix, showing that astrocytes presented the characteristic star-shaped morphology in the hydrogel. g) Permeability coefficients (*P_app_*), calculated from the diffusion of 4 kDa FITC-dextran through a membrane with just hydrogel (No cell), an endothelial monolayer with hydrogel (Monoculture), an endothelial monolayer cocultured with HAs and HPs (Coculture), with and without the application of gravity-driven medium flow (n = 6 for each condition); h) Gene expression of ECs in monoculture and coculture under static and dynamic culturing conditions (n = 6 for each condition, all the data were normalized to the TFRC in the coculture system under dynamic conditions); i) Normalized TEER values were measured using the integrated sensor every hour in the coculture system (the graph shows the mean values ± standard deviation in the form of a shaded envelope of the normalized TEER values at 1 kHz, n = 6). Data represent mean values ± s.d., \**p* ≤ 0.05, \*\**p* ≤ 0.01; \*\*\**p* ≤ 0.001; \*\*\*\**p* ≤ 0.0001; ns, not significant. Scale bars = 50 μm.

We then studied BBB tightness and EC monolayer formation under different culture conditions, namely dynamic (i.e., by perfusing via asymmetric, periodic tilting of the platform) and static culturing (i.e., by maintaining the platform in a horizontal configuration), and in monocultures of ECs or triple cocultures of ECs, HPs, and HAs. Barrier formation was characterized using real-time TEER, end-point permeability measurements, immunofluorescence staining and protein expression via RT-qPCR (Fig. 3b). For all conditions, the ECs were cultured on chip during 4 days before characterization. Under dynamic culturing, the ECs showed an elongated morphology and orientation in the flow direction, with a higher localization of the tight-junction protein zona occludens 1 (ZO-1) and vascular endothelial cadherin (VE-CAD) at the cell-to-cell junctions with respect to ECs under static conditions (Fig. 3c). Coculturing with HAs and HPs further promoted the localization of ZO-1 and VE-CAD, an indication of tighter barrier formation.[44, 45] Confocal microscopy imaging of the BBB structure confirmed that, under coculture conditions, the HPs grew on the opposite side of the porous membrane (Fig. 3d, e) in close proximity to the EC monolayer, while HAs exhibited characteristic star-shaped morphologies with extended end feet in the hydrogel matrix (Fig. 3d, f and Supplementary Fig. 7). These findings indicate that the BBB model featured an organized 3D multilayer cellular structure.

We assessed the barrier permeability by measuring the paracellular transport of 4 kDa FITC-Dextran[17, 18] under the different culturing conditions (Fig. 3g). The BBB models cultured under static conditions, either as EC monoculture or triple coculture, did not show significant differences in permeability (*P_app_* =24.8 ± 1.4 × 10^−7^ cm/s and *P_app_* =17.3 ± 1.1 × 10^−7^ cm/s, for monoculture and coculture conditions, respectively). However, their permeability was lower than that of the acellular control group (*P_app_* = 38.6 ± 5.5 × 10^−7^ cm/s), which suggests that a cellular barrier had formed on chip. Exposure to shear stress further decreased the permeability of the BBB models, which resulted in a 2-fold decrease for the monoculture BBB model (*P_app_* = 10.5 ± 0.5 × 10^−7^ cm/s) and a 12-fold decrease for the coculture model (*P_app_* of 1.4 ± 0.3 × 10^−7^ cm/s). The application of shear stress, induced by the gravity-driven flow, seemingly promoted the formation of a tighter barrier layer, which confirms the importance of the presence of shear stress in *in vitro* barrier models.[32, 46] Furthermore, the permeability of the coculture barrier model was ~7-fold lower than that of the monoculture barrier with applied shear stress, which indicated that the coculture of ECs with HPs and HAs enhanced the tightness of the BBB model. Finally, the *P_app_* of the dynamic, coculture BBB model was comparable to previously reported permeability values for both, *in vitro* (*P_app_* ~ 10^−6^ ~ 10^−7^ cm/s)[18, 47–50] and *in vivo* (~ 4 × 10^−7^ cm/s in rat)[51, 52] BBB studies, which confirms that the quasi-unidirectional, tilting-induced medium flow effectively supported the formation of a tight barrier on chip.

Next, we performed RT-qPCR to determine the expression of ZO-1 and VE-CAD, as markers of tight-junction-protein expression and cellular polarity (Fig. 3h).[18, 53] ECs in the coculture BBB models showed a lower expression of these proteins in comparison to their monoculture counterparts under static and dynamic conditions. This effect is particularly evident for the static coculture condition, which showed a much lower protein expression than the corresponding monoculture model. No significant difference in ZO-1 and VE-CAD expression was measured in EC monocultures under static or dynamic conditions. These measurements show that gene expression alone is not a good indicator of barrier tightness, as higher ZO-1 or VE-CAD gene expressions are seemingly not correlated to tighter cellular barriers[5, 54–56].

Lastly, the BBB formation of the coculture model under dynamic conditions was continuously monitored by using the integrated TEER sensor. The trans-barrier resistance was recorded at multiple frequencies to improve the reliability of the detection (Supplementary Fig. 8). To compare different chips, the TEER values were normalized between 0% (TEER at day 0, when ECs were loaded into the chip) and 100% at day 4 (Fig. 3i). The normalization was necessary as different chips featured different baseline values owing to small variations in the alignment of the electrodes with the fluidic network during fabrication. The continuous TEER measurements enabled us to follow and confirm the formation of a tight cellular barrier on chip.

### Monitoring rapid disruption of the BBB

We then sought to verify whether our coculture system could be used to monitor rapid variations in the BBB structure. To this end, we exposed the BBB to ethylenediaminetetraacetate (EDTA), a strong chelator of divalent cations, such as calcium (Ca^2+^) and magnesium (Mg^2+^), to induce cation depletion in the EC microchannel, which affects barrier tightness and promotes cell detachment.[36, 57–59] To compare quantitative results obtained by TEER measurements with morphological and functional changes in barrier integrity, we simultaneously monitored focal adhesion and cell permeability disruption by high-resolution time-lapse microscopy. Barrier permeability was measured after EDTA treatment using the FITC-Dextran fluorescent tracer (Fig. 4a).

**Fig. 4.**
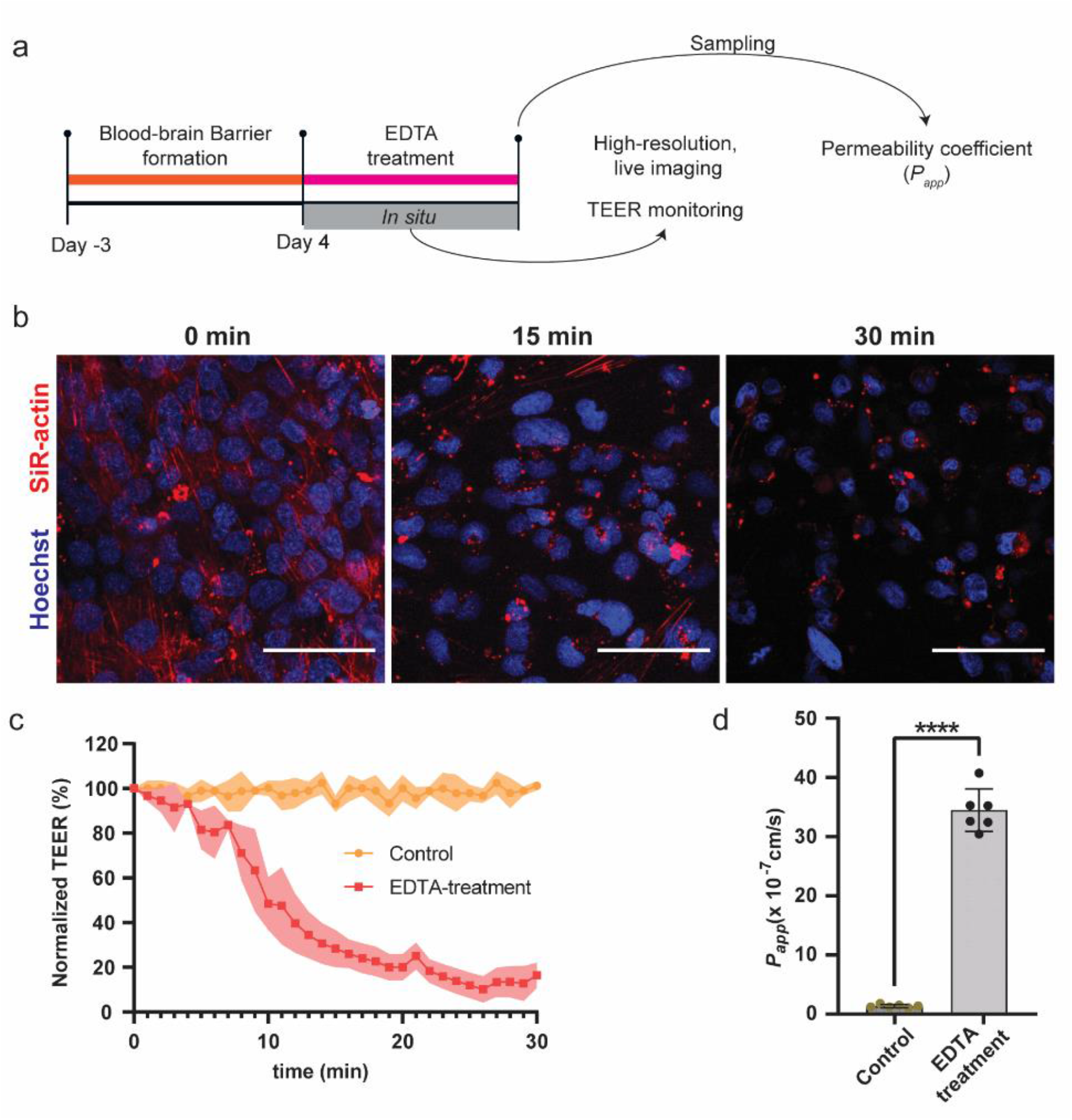
a) Schematic of the experimental workflow for monitoring EDTA-induced barrier disruption; b) High-resolution, live imaging of ECs on chip. The endothelial layer was stained for F-actin (SiR-actin, red) and cell nuclei (Hoechst, blue). EDTA induced actin-fiber aggregation within 15 min and cell detachment of the barrier within 30 min of the treatment; c) TEER values were recorded every 2 min after EDTA dosage (the graph shows mean values ± standard deviation in the form of a shaded envelope of the normalized TEER values at 1 kHz, n = 6 for each condition); d) Permeability measurements were acquired by measuring the diffusion of 4-kDa FITC-dextran after 30 min of EDTA treatment (n = 6 for each condition); Control group without EDTA treatment. Data represent mean values ± s.d., \**p* ≤ 0.05, \*\**p* ≤ 0.01; \*\*\**p* ≤ 0.001; \*\*\*\**p* ≤ 0.0001. Scale bars = 50 μm.

Live imaging was performed by staining the cellular nuclei with Hoechst and the actin cytoskeleton with SiR-actin. Within 15 minutes of exposure to EDTA, we observed the collapse of the actin filaments in the ECs, which indicates a loss of focal contacts of the ECs to the membrane support and a drastic variation in cell morphology,[60] while cell detachment was observed after 30 minutes (Fig. 4b). In parallel, the TEER value started to decrease as the EDTA solution was injected into the microchannel, showing a TEER decrease of 71.7 ± 8.3% of the initial TEER value within the first 15 minutes of EDTA exposure (Fig. 4c). The rate of TEER decrease then reduced, and the TEER value stabilized at ~12% of the original TEER value after ~30 minutes. The pronounced drop of TEER upon EDTA application is in line with previously reported measurements in other organ-on-chip models.[36, 57] Finally, we assessed the barrier permeability after 30 minutes of EDTA treatment to confirm these data. The *P_app_* of the 4 kDa Dextran- FITC was significantly increased, showing a ~25-fold increase after EDTA treatment (*P_app_* = 34.8 ± 3.8 × 10^−7^ cm/s) compared to the control condition (*P_app_* = 1.4 ± 0.3 × 10^−7^ cm/s; Fig. 4d). These results demonstrate that our BBB platform is capable of real-time monitoring - through TEER and high-resolution microscopy - rapid variations in the *in vitro* BBB model upon external insults.

### The BBB platform as an effective *in vitro* model for recapitulating ischemic injuries

After confirming that our system was capable of monitoring rapid variations in BBB permeability and EC-layer organization, we challenged our barrier model with an ischemia-like insult by exposing the model to oxygen-glucose-deprivation (OGD), which is a commonly applied method to induce ischemic injuries *in vitro*[61] (Fig. 5a). OGD conditions were applied by replacing the EAPM by glucose-free DMEM and by flushing the culture chamber with 95% N_2_ - 5% CO_2_.

**Fig. 5.**
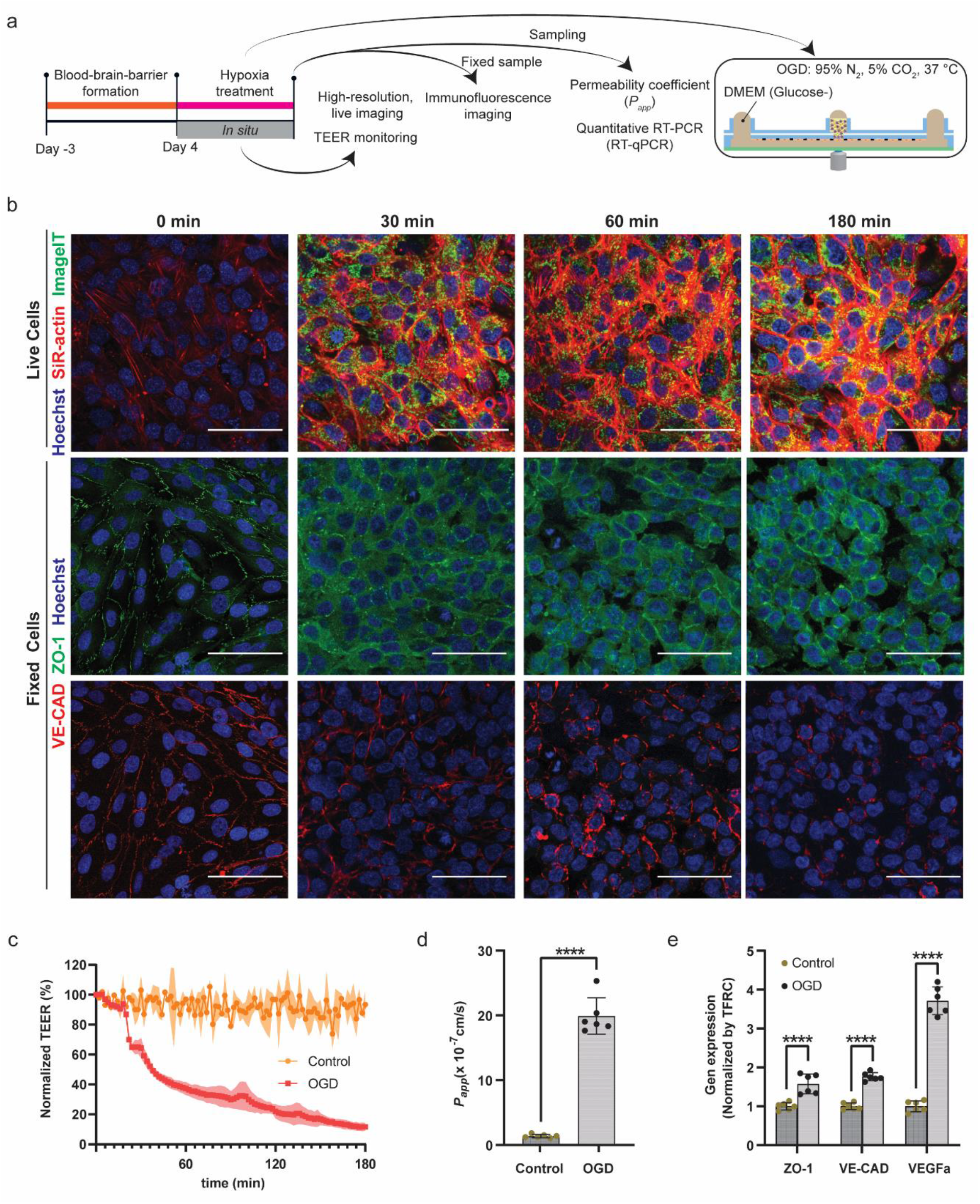
a) Schematic of the experimental workflow for recapitulating ischemic injury on chip; b) The cellular barrier on chips was imaged continuously, and separate chips were fixed for immunofluorescence image analysis after 0 min, 30 min, 1 h, and 3 h of OGD exposure. For live imaging, cells were stained for F-actin (SiR-actin, red), hypoxia (Image-IT, green) and cell nuclei (Hoechst, blue). Large actin stress-fiber aggregates appeared after 30 minutes of OGD exposure. After 1 h, cells started to assume a round morphology, which compromised barrier tightness. The effect was even more visible after 3h of OGD insult. For fixed, immunofluorescent-stained images, the EC layer was labeled for ZO-1 (green), VE-CAD (red) and cell nuclei (blue). After 30 minutes of OGD, ZO-1 and VE-CAD showed less localization at cell boundaries with respect to the initial barrier; at later time points, the barrier showed a drastic decrease in localization of the adherent and junction proteins, as well as a round cell morphology and openings and voids; c) TEER values were acquired every 2 min from chips exposed to hypoxia and control conditions after the OGD had been applied; (graph shows mean values ± s.d. in the form of a shaded envelope of the normalized TEER values at 1 kHz, n = 6 for each condition); d) Permeability measurements were carried out after 3 h of OGD exposure by measuring the diffusion of 4 kDa FITC-dextran across the barrier (n = 6 for each condition); e) Gene expression of ECs under normal and OGD culture conditions, including the ZO-1 tight junction protein, the VE-CAD adherent junction protein and the vascular growth factors VEGFa; n = 6 for each condition). Control group without hypoxia treatment. Data represent mean values ± s.d., \**p* ≤ 0.05, \*\**p* ≤ 0.01; \*\*\**p* ≤ 0.001; \*\*\*\**p* ≤ 0.000. Scale bars: 50 μm.

We monitored the barrier behavior by continuously recording the TEER value and by using high-resolution microscopy, for which we used live fluorescence markers for actin (SiR-actin), cell nuclei (Hoechst) and cellular hypoxia (Image-IT). Within 4 minutes of incubation under OGD conditions, the fluorescence signal of the hypoxia indicator appeared in the cellular barrier (Supplementary Fig. 9), which indicated that our open-microfluidic solution enabled us to quickly expose the BBB to hypoxia stress conditions. Concurrently, the cells started to show a marked increase in actin polymerization, as evidenced by the increased fluorescence of the actin filaments through the SiR-actin staining. A strong SiR-actin signal appeared within 30 minutes of OGD exposure, and actin stress fibers were clearly visible by using live, confocal microscopy. After 60 min of OGD exposure, the actin stress fibers started to clump together, which led to the contraction of the ECs, as clearly visible at 180 minutes post-OGD exposure (Fig. 5b and Supplementary Video 1). The behavior of the ECs in our BBB devices confirmed that actin stress-fiber polymerization caused an increase in cytoskeletal tension of the ECs, which led to an impairment of the junctions between ECs and to a hyperpermeability of the barrier.[15]

The cellular barriers were also fixed at different time points to inspect the barrier reorganization through immunofluorescence staining (Fig. 5b). Within 30 min of OGD exposure, although the barrier remained intact, ZO-1 and VE-CAD showed less localization at cellular junctions, which indicated that the junction proteins quickly rearranged under OGD. After 60 min of OGD exposure, the cellular barrier showed clear signs of disruption, and the ECs started assuming a rounder morphology. After 180 minutes of OGD exposure, the barrier was completely compromised, showing holes and void spaces between the cells.

The continuous TEER measurements showed that the TEER value started decreasing 6 minutes after exposure to OGD, indicating hypoxia-stress-induced barrier dysfunction before actin polymerization could be observed through live imaging (Fig. 5c, Supplementary Fig. 9). TEER decreased by 6.7 ± 1.8% within the first 10 min, and, then, the value rapidly dropped by 35.7 ± 6.7% within 30 minutes of exposure. Eventually, the TEER value dropped by 88.4 ± 2.4% within 3 hours of the OGD insult, which indicated the disruption of the barrier on chip. We subsequently measured the permeability of the barrier using the 4 kDa dextran-FITC tracer to further evaluate the tightness of the barrier on the chips (Fig. 5d). In accordance with the TEER measurements, the *P_app_* was significantly increased by ~20 fold after 3 h of OGD exposure (22.2 ± 4.4 × 10^−7^ cm/s) compared to the control condition (1.2 ± 0.1 × 10^−7^ cm/s). The TEER and permeability measurements confirmed that the barrier was disrupted by the OGD insult.

RT-qPCR analysis of the harvested ECs after 3 hours of exposure to OGD conditions showed that OGD led to significantly increased expression of ZO-1 (1.57 ± 0.25), VE-CAD (1.76 ± 0.10), and VEGFa (3.71 ± 0.36) compared to normoxic control samples (Fig. 5e). These results suggest a compensatory mechanism for the reduction of tight junctions to counteract barrier disruption.

## Discussion

In this work, we described the development and validation of a microfluidic platform with an integrated transparent TEER sensor to model the BBB *in vitro*. The platform has the following advantages over previously reported *in vitro* BBB models: (1) it combines a 2D endothelial monolayer with a 3D brain microenvironment to reconstruct the BBB structure on chip; (2) it features a tubing- and pump-free fluidic system to induce flow across the endothelial layer, which enables straightforward parallelization and provides increased throughput; (3) it features a quasi-unidirectional, gravity-driven flow that provides shear stress at physiological levels; (4) it includes an open-microfluidic network for simple liquid exchange and sampling; (5) it contains integrated TEER electrodes, which were optimized for uniform current density across the cellular layer; (6) the integrated ITO electrodes are fully transparent and enable simultaneous high-resolution imaging and TEER measurements.

TEER measurements are a standard, non-invasive method for evaluating barrier tightness in *in vitro* models, and both, integrated[23, 36, 38] or plug-in wire electrodes[13, 18, 20, 48, 62] have been used in literature to monitor the characteristics of microfluidics-based BBB models. Wire electrodes can be easily integrated or inserted on demand within microfluidic systems, without requiring specialized and complex fabrication steps. However, this type of electrode usually does not provide uniform current density across barrier layers due to the wire geometry. For non-integrated electrodes, differences in electrode placement between measurements add an additional source of variation to the TEER measurements. Moreover, plug-in wire-electrodes, which are typically inserted into the inlet and outlet of the microfluidic device and distant to the cellular barrier, may lead to strong artifacts in TEER measurements due to resistance contributions of the cell culture medium in the microfluidic system containing the electrodes and the cellular barrier.[35, 38, 63] Therefore, the sensitivity and the reliability of TEER measurements and the corresponding information on the tightness of the whole barrier may be compromised.[35] To obtain sensitive TEER measurements, thin-film TEER sensors have been integrated within microfluidic-based systems[23, 36, 64] using different electrode materials (e.g., Au, Ag, Pt) on a variety of substrates (e.g., glass, PET, PC). However, the low transparency of these thin-film electrodes prevents direct optical access to the cellular barriers, and imaging can only be performed as an end-point examination, after disassembling of the platform. Alternatively, the electrodes do not extend across the whole barrier region, which results in non-uniform current densities across the barrier layer. Here, we overcame such limitations by integrating transparent, thin-film TEER sensing electrodes in close proximity to and on both sides of the cellular barrier. This approach enabled to reduce the contribution of the electrical resistance of the cell-culture medium and helped to avoid measurement fluctuations as a consequence of varying sensor placements during multi-day measurements. Through FEM simulations, we optimized the design of the electrodes so as to obtain a uniform current density across the barrier layer. The electrode geometry was designed to enable the use of open-microfluidic structures to provide direct access to the “brain” side of the BBB model for simple loading of the hydrogel matrix for 3D cell culturing and for the sampling of the supernatant. We found that a spiral-shaped electrode, in combination with a ring electrode for the microfluidic open access, provided uniform current density across the barrier layer. Although the electrodes were located within the optical path to the barrier layer, the high transparency of the ITO electrode and the use of a microscope coverslip as substrate material ensured compatibility with live, high-resolution, confocal microscopy, so that rapid variations in the organization of the cellular barrier layer could be observed with high spatiotemporal resolution.

In a first step in the realization of our *in vitro* BBB model, we confirmed that a common medium formulation could be used for coculturing ECs, HAs, and HPs in the platform. Our results showed that all cell types had a viability of more than 90% in the EAPM coculture medium, with ECs even presenting higher viability than in their standard medium despite the removal of VEGF to improve barrier tightness. We reproduced the BBB arrangement on chip by combining a 2D endothelial monolayer with a 3D brain-like culture microenvironment for the HAs and HPs. *In vivo,* HPs surround the endothelial microvessels and fundamentally affect the function of the BBB in combination with the HAs.[65, 66] In our model, the HPs were cultured in close proximity to the EC layer and within the same hydrogel compartments as the HAs to support the structure and function of the BBB cellular barrier on chip. The selected hydrogel matrix for culturing of HAs in 3D supported the growth of HAs that displayed long branches and end feet in close contact with the HPs (Supplementary Fig. 7), which contributed to the formation of a robust and functional BBB on chip.[18, 67]

In vessels, blood flow generates shear stress on the endothelial cell layer, which is known to maintain the structure and function of the vessel barrier layer by regulating endothelial cytoskeleton and vascular homeostasis.[32, 68]. To recapitulate this effect *in vitro*, several BBB models have relied on external pumping systems to generate medium flow inside microfluidic devices and to improve the tightness of the barrier model.[18, 20, 21, 47, 62, 69] However, tubing and pump systems impede parallelization for medium-to-high-throughput studies due to the complex experimental setup that is required to operate each BBB model.[17] Here, the medium flow was achieved by tilting the platform[70, 71] without the use of external tubing and pumping devices. Although the generated flow was not perfectly unidirectional, as would have been the case for using an external pump, our results show that the asymmetric tilting profile and microfluidic-chip geometry enabled to generate shear stress with a preferential directionality and within the physiological range, which improved the integrity and functionality of the *in vitro* BBB model. Our permeability measurements and immunofluorescence images show that the BBB, cultured under perfusion conditions, formed a tighter barrier, with localized junction and adherent proteins between ECs in the barrier, in comparison to static culture conditions. Moreover, ECs showed an improved orientation along the flow direction, while ECs in static conditions presented a round morphology.

In the monoculture system, we did not see a significant difference in the expression of junction-specific proteins between conditions with and without shear stress, however, the permeability of the barrier was significantly lower under dynamic culturing conditions compared to static culturing. These observations may indicate that the shear stress does not affect the expression of junction-specific proteins at the mRNA level, but promotes the specific localization of such proteins, as has been previously reported for human iPSC-derived endothelial cells.[72] In the coculture system on our chips, the expression of these junction-specific proteins was slightly downregulated under perfusion conditions in comparison to the monoculture on chip. However, the dynamic, coculture BBB model exhibited a significantly lower permeability, which evidences the importance of coculturing ECs with HAs and HPs and of using a dynamic culturing environment to promote junction-protein localization at cell-cell boundaries in the EC layer to achieve the formation of a tight barrier.[73] The formation of a tight on-chip barrier was further confirmed by the continuous increase in TEER value during EC culturing, which was detected by the integrated sensor.

After validating the BBB model, we investigated whether the integrated transparent TEER sensor could be used to monitor rapid variations in the cellular barrier upon exposure to external insults, concurrently with high-resolution microscopy. To this end, we exposed the BBB model to EDTA, which is known to disrupt the junction protein assemblies and promote cell detachment.[57, 59] The EDTA exposure caused a rapid breakage of the barrier, which, in turn, resulted in a rapid drop in the TEER values, while morphological changes and loss of focal adhesion of the ECs, as well as cellular detachment from the porous membrane, were detected via high-resolution microscopy. The EDTA test confirmed that the integrated TEER sensor was able to detect and quantify a rapid variation in barrier integrity, which could be simultaneously confirmed through live, confocal microscopy.

Finally, we used the BBB platform to recapitulate the disruption of the BBB on chip by an ischemic insult upon applying oxygen-glucose-deprivation conditions. The TEER values rapidly decreased within the first few minutes after OGD onset. The OGD triggered actin polymerization in the ECs to induce cytoskeletal alteration, which is known to be an initiator of BBB rupture.[15, 74] After ~30 minutes, live imaging revealed that actin stress fibers were continuously being formed within the EC layer, which produced contractile forces in the cells, with the actin cytoskeleton directly pulling on the adherent and tight junction proteins VE-CAD and ZO-1.[74] Gene-expression analysis revealed that these adherent and junction proteins were highly expressed under hypoxia conditions in comparison to the normoxia samples, which may indicate that the EC layer tries to maintain its integrity by promoting the expression of the cellular junctional protein to counteract the cytoskeletal contractile forces. Live and immunofluorescence microscopy images showed that the ECs ultimately contracted, and VE-CAD and ZO-1 were removed from the cell-cell contact boundaries and internalized. Our model also confirmed the over-expression of VEGFa due to hypoxic stress, which has been hypothesized being a defense mechanism of the BBB during ischemia, through which the vessels promote angiogenesis to respond to barrier-induced OGD stress.[75, 76] TEER value decreases were evident before the appearance of the large stress-fiber assemblies and cell contractions, which indicates that a leaky barrier had formed before drastic morphological changes in the EC layer occurred. This finding demonstrates the high sensitivity of TEER as a real-time monitoring method for barrier tightness and the advantages of performing concurrent TEER and microscopy measurements to investigate barrier-disruption events.

## Conclusion

In summary, we established a robust, highly integrated, and user-friendly BBB model platform that can be used to recapitulate barrier functions and to investigate rapid variations of the endothelial layer in response to external insults. Using the BBB platform to mimic cerebral ischemia, the model can contribute to elucidating the mechanism involved in barrier disruption, which will facilitate the discovery of BBB stabilizers to reduce brain edema and neurological damage. Additionally, the developed BBB platform provides a tool for the development of therapeutic agents for brain and BBB-related diseases. The high level of tightness of the BBB renders the developed system a suitable *in vitro* tool for screening novel therapeutic compounds and strategies for BBB crossing, a major roadblock in the development of compounds for CNS diseases. Moreover, the combination of an open-microfluidic, hanging-drop BBB platform will provide the possibility of easily integrating preformed brain-disease tissue models into the system to investigate both, the barrier permeability to candidate compounds as well as compound efficacy on the target tissue.

## Methods

All experimental procedures and methods are provided in the Supplementary Information.

## Data availability

The main data supporting the results in this study are available within the paper and its Supplementary Information. The raw and analysed datasets generated during the study available for research purposes from the corresponding author on reasonable request.

## Code availability

MATLAB was used to calculate the flow rate through the microchannel during the experiments. The codes used for the calculation can be provided on request.

## Acknowledgments

WW was supported by the China Scholarship Council (grant no. 201804910890). The authors acknowledge the clean-room facility and single-cell facility at D-BSSE, ETH Zürich, for help and support. Further, the authors would like to thank Dr. D. Eletto, ETH Zürich, for helping with the RT-qPCR. The authors thank Dr. C. Campa from the Italian Institute of Genomic Medicine for valuable inputs and comments to the manuscript.

## Author contributions

WW, AH and MMM conceived and designed the research. WW designed and developed the TEER sensor under the supervision of FC and MMM. WW designed and developed the microfluidic platform, performed experiments, and performed the data analysis under the supervision of MMM. All authors wrote the manuscript and approved the final manuscript.

## References

[1] A. Oddo, B. Peng, Z. Tong, Y. Wei, W.Y. Tong, H. Thissen, N.H. Voelcker, Advances in Microfluidic Blood-Brain Barrier (BBB) Models, Trends Biotechnol (2019).

[2] A. Bhalerao, F. Sivandzade, S.R. Archie, E.A. Chowdhury, B. Noorani, L. Cucullo, In vitro modeling of the neurovascular unit: advances in the field, Fluids Barriers CNS 17(1) (2020) 22.

[3] J. Stockwell, N. Abdi, X. Lu, O. Maheshwari, C. Taghibiglou, Novel central nervous system drug delivery systems, Chem Biol Drug Des 83(5) (2014) 507–20.

[4] K. Boye, L.H. Geraldo, J. Furtado, L. Pibouin-Fragner, M. Poulet, D. Kim, B. Nelson, Y. Xu, L. Jacob, N. Maissa, D. Agalliu, L. Claesson-Welsh, S.L. Ackerman, A. Eichmann, Endothelial Unc5B controls blood-brain barrier integrity, Nat Commun 13(1) (2022) 1169.

[5] A. Gerhartl, N. Pracser, A. Vladetic, S. Hendrikx, H.P. Friedl, W. Neuhaus, The pivotal role of micro-environmental cells in a human blood-brain barrier in vitro model of cerebral ischemia: functional and transcriptomic analysis, Fluids Barriers CNS 17(1) (2020) 19.

[6] H. Kadry, B. Noorani, L. Cucullo, A blood–brain barrier overview on structure, function, impairment, and biomarkers of integrity, Fluids and Barriers of the CNS 17(1) (2020).

[7] A. Tunon-Ortiz, T.J. Lamb, Blood brain barrier disruption in cerebral malaria: Beyond endothelial cell activation, PLoS Pathog 15(6) (2019) e1007786.

[8] R. Khatri;, A.M. McKinney;, B. Swenson;, V. Janardhan, Blood–brain barrier, reperfusion injury, and hemorrhagic transformation in acute ischemic stroke, Neurology 79(13) (2012) 6.

[9] A.D. Wong, M. Ye, A.F. Levy, J.D. Rothstein, D.E. Bergles, P.C. Searson, The blood-brain barrier: an engineering perspective, Front Neuroeng 6 (2013) 7.

[10] R. Cecchelli, V. Berezowski, S. Lundquist, M. Culot, M. Renftel, M.P. Dehouck, L. Fenart, Modelling of the blood-brain barrier in drug discovery and development, Nat Rev Drug Discov 6(8) (2007) 650–61.

[11] G.C. Terstappen, A.H. Meyer, R.D. Bell, W. Zhang, Strategies for delivering therapeutics across the blood-brain barrier, Nat Rev Drug Discov 20(5) (2021) 362–383.

[12] N. Zhao, Z. Guo, S. Kulkarni, D. Norman, S. Zhang, T.D. Chung, R.F. Nerenberg, R.M. Linville, P. Searson, Engineering the Human Blood-Brain Barrier at the Capillary Scale using a Double-Templating Technique, Advanced Functional Materials (2022).

[13] J.J. Jamieson, R.M. Linville, Y.Y. Ding, S. Gerecht, P.C. Searson, Role of iPSC-derived pericytes on barrier function of iPSC-derived brain microvascular endothelial cells in 2D and 3D, Fluids and Barriers of the CNS 16(1) (2019).

[14] C. Kulczar, K.E. Lubin, S. Lefebvre, D.W. Miller, G.T. Knipp, Development of a direct contact astrocyte-human cerebral microvessel endothelial cells blood-brain barrier coculture model, J Pharm Pharmacol 69(12) (2017) 1684–1696.

[15] Y. Shi, L. Zhang, H. Pu, L. Mao, X. Hu, X. Jiang, N. Xu, R.A. Stetler, F. Zhang, X. Liu, R.K. Leak, R.F. Keep, X. Ji, J. Chen, Rapid endothelial cytoskeletal reorganization enables early blood-brain barrier disruption and long-term ischaemic reperfusion brain injury, Nat Commun 7 (2016) 10523.

[16] K. Hatherell, P.O. Couraud, I.A. Romero, B. Weksler, G.J. Pilkington, Development of a three-dimensional, all-human in vitro model of the blood-brain barrier using mono-, co-, and tri-cultivation Transwell models, J Neurosci Methods 199(2) (2011) 223–9.

[17] J. Kim, K.T. Lee, J.S. Lee, J. Shin, B. Cui, K. Yang, Y.S. Choi, N. Choi, S.H. Lee, J.H. Lee, Y.S. Bahn, S.W. Cho, Fungal brain infection modelled in a human-neurovascular-unit-on-a-chip with a functional blood-brain barrier, Nat Biomed Eng 5(8) (2021) 830–846.

[18] S.I. Ahn, Y.J. Sei, H.J. Park, J. Kim, Y. Ryu, J.J. Choi, H.J. Sung, T.J. MacDonald, A.I. Levey, Y. Kim, Microengineered human blood-brain barrier platform for understanding nanoparticle transport mechanisms, Nat Commun 11(1) (2020) 175.

[19] B.M. Maoz, A. Herland, E.A. FitzGerald, T. Grevesse, C. Vidoudez, A.R. Pacheco, S.P. Sheehy, T.E. Park, S. Dauth, R. Mannix, N. Budnik, K. Shores, A. Cho, J.C. Nawroth, D. Segre, B. Budnik, D.E. Ingber, K.K. Parker, A linked organ-on-chip model of the human neurovascular unit reveals the metabolic coupling of endothelial and neuronal cells, Nat Biotechnol 36(9) (2018) 865–874.

[20] T.-E. Park, N. Mustafaoglu, A. Herland, R. Hasselkus, R. Mannix, E.A. FitzGerald, R. Prantil-Baun, A. Watters, O. Henry, M. Benz, H. Sanchez, H.J. McCrea, L.C. Goumnerova, H.W. Song, S.P. Palecek, E. Shusta, D.E. Ingber, Hypoxia-enhanced Blood-Brain Barrier Chip recapitulates human barrier function and shuttling of drugs and antibodies, Nature Communications 10(1) (2019).

[21] Y. Shin, S.H. Choi, E. Kim, E. Bylykbashi, J.A. Kim, S. Chung, D.Y. Kim, R.D. Kamm, R.E. Tanzi, Blood–Brain Barrier Dysfunction in a 3D In Vitro Model of Alzheimer’s Disease, Advanced Science (2019).

[22] P. Motallebnejad, A. Thomas, S.L. Swisher, S.M. Azarin, An isogenic hiPSC-derived BBB-on-a-chip, Biomicrofluidics 13(6) (2019) 064119.

[23] R. Booth, H. Kim, Characterization of a microfluidic in vitro model of the blood-brain barrier (muBBB), Lab Chip 12(10) (2012) 1784–92.

[24] B. Zhang, A. Korolj, B.F.L. Lai, M. Radisic, Advances in organ-on-a-chip engineering, Nature Reviews Materials 3(8) (2018) 257–278.

[25] Y. Xia, J.C. Izpisua Belmonte, Design Approaches for Generating Organ Constructs, Cell Stem Cell 24(6) (2019) 877–894.

[26] S.N. Bhatia, D.E. Ingber, Microfluidic organs-on-chips, Nat Biotechnol 32(8) (2014) 760–72.

[27] Z. Lyu, J. Park, K.M. Kim, H.J. Jin, H. Wu, J. Rajadas, D.H. Kim, G.K. Steinberg, W. Lee, A neurovascular-unit-on-a-chip for the evaluation of the restorative potential of stem cell therapies for ischaemic stroke, Nat Biomed Eng 5(8) (2021) 847–863.

[28] S. Chien, Effects of disturbed flow on endothelial cells, Ann Biomed Eng 36(4) (2008) 554–62.

[29] J.J. Chiu, S. Chien, Effects of disturbed flow on vascular endothelium: pathophysiological basis and clinical perspectives, Physiol Rev 91(1) (2011) 327–87.

[30] O.F. Khan, M.V. Sefton, Endothelial cell behaviour within a microfluidic mimic of the flow channels of a modular tissue engineered construct, Biomed Microdevices 13(1) (2011) 69–87.

[31] F. Garcia-Polite, J. Martorell, P. Del Rey-Puech, P. Melgar-Lesmes, C.C. O’Brien, J. Roquer, A. Ois, A. Principe, E.R. Edelman, M. Balcells, Pulsatility and high shear stress deteriorate barrier phenotype in brain microvascular endothelium, J Cereb Blood Flow Metab 37(7) (2017) 2614–2625.

[32] L. Cucullo, M. Hossain, V. Puvenna, N. Marchi, D. Janigro, The role of shear stress in Blood-Brain Barrier endothelial physiology, BMC Neurosci 12 (2011) 40.

[33] E. Lauranzano, E. Campo, M. Rasile, R. Molteni, M. Pizzocri, L. Passoni, L. Bello, D. Pozzi, R. Pardi, M. Matteoli, A. Ruiz-Moreno, A Microfluidic Human Model of Blood– Brain Barrier Employing Primary Human Astrocytes, Advanced Biosystems (2019).

[34] T.-E. Park, N. Mustafaoglu, A. Herland, R.M. Hasselkus, R. Mannix, E.A. FitzGerald, R. Prantil-Baun, A. Watters, O. Henry, M. Benz, H. Sanchez, H.J. McCrea, L. Christova Goumnerova, H.W. Song, S.P. Palecek, E. Shusta, D.E. Ingber, Hypoxia-enhanced Blood-Brain Barrier Chip recapitulates human barrier function, drug penetration, and antibody shuttling properties, (2018).

[35] B. Srinivasan, A.R. Kolli, M.B. Esch, H.E. Abaci, M.L. Shuler, J.J. Hickman, TEER measurement techniques for in vitro barrier model systems, J Lab Autom 20(2) (2015) 107–26.

[36] O.Y.F. Henry, R. Villenave, M.J. Cronce, W.D. Leineweber, M.A. Benz, D.E. Ingber, Organs-on-chips with integrated electrodes for trans-epithelial electrical resistance (TEER) measurements of human epithelial barrier function, Lab Chip 17(13) (2017) 2264–2271.

[37] B. Srinivasan, A.R. Kolli, Transepithelial/Transendothelial Electrical Resistance (TEER) to Measure the Integrity of Blood-Brain Barrier, Blood-Brain Barrier2019, pp. 99–114.

[38] M.W. van der Helm, O.Y.F. Henry, A. Bein, T. Hamkins-Indik, M.J. Cronce, W.D. Leineweber, M. Odijk, A.D. van der Meer, J.C.T. Eijkel, D.E. Ingber, A. van den Berg, L.I. Segerink, Non-invasive sensing of transepithelial barrier function and tissue differentiation in organs-on-chips using impedance spectroscopy, Lab Chip (2019).

[39] K.A. Kim, D. Kim, J.H. Kim, Y.J. Shin, E.S. Kim, M. Akram, E.H. Kim, A. Majid, S.H. Baek, O.N. Bae, Autophagy-mediated occludin degradation contributes to blood-brain barrier disruption during ischemia in bEnd.3 brain endothelial cells and rat ischemic stroke models, Fluids Barriers CNS 17(1) (2020) 21.

[40] A. W, T. D, R. Pt, Blood-brain barrier dysfunction in ischemic stroke: targeting tight junctions and transporters for vascular protection, Am J Physiol Cell Physiol. 315(3) (2018) 16.

[41] S. Page, A. Munsell, A.J. Al-Ahmad, Cerebral hypoxia/ischemia selectively disrupts tight junctions complexes in stem cell-derived human brain microvascular endothelial cells, Fluids Barriers CNS 13(1) (2016) 16.

[42] P. Silacci, A. Desgeorges, L. Mazzolai, C. Chambaz, D. Hayoz, Flow Pulsatility Is a Critical Determinant of Oxidative Stress in Endothelial Cells, Hypertension 38(5) (2001) 4.

[43] C.F. D. Jr., S.R. Bussolari, J. M. A. Gimbrone, P.F. Davies, The Dynamic Response of Vascular Endothelial Cells to Fluid Shear Stress, J Biomech Eng 103 (1981) 8.

[44] E. Sasson, S. Anzi, B. Bell, O. Yakovian, M. Zorsky, U. Deutsch, B. Engelhardt, E. Sherman, G. Vatine, R. Dzikowski, A. Ben-Zvi, Nano-scale architecture of blood-brain barrier tight-junctions, Elife 10 (2021).

[45] T. Koto, K. Takubo, S. Ishida, H. Shinoda, M. Inoue, K. Tsubota, Y. Okada, E. Ikeda, Hypoxia disrupts the barrier function of neural blood vessels through changes in the expression of claudin-5 in endothelial cells, Am J Pathol 170(4) (2007) 1389–97.

[46] M.W. van der Helm, A.D. van der Meer, J.C. Eijkel, A. van den Berg, L.I. Segerink, Microfluidic organ-on-chip technology for blood-brain barrier research, Tissue Barriers 4(1) (2016) e1142493.

[47] R.M. Linville, J.G. DeStefano, M.B. Sklar, Z. Xu, A.M. Farrell, M.I. Bogorad, C. Chu, P. Walczak, L. Cheng, V. Mahairaki, K.A. Whartenby, P.A. Calabresi, P.C. Searson, Human iPSC-derived blood-brain barrier microvessels: validation of barrier function and endothelial cell behavior, Biomaterials 190-191 (2019) 24–37.

[48] Y.I. Wang, H.E. Abaci, M.L. Shuler, Microfluidic blood-brain barrier model provides in vivo-like barrier properties for drug permeability screening, Biotechnol Bioeng 114(1) (2017) 184–194.

[49] J.G. DeStefano, J.J. Jamieson, R.M. Linville, P.C. Searson, Benchmarking in vitro tissue-engineered blood-brain barrier models, Fluids Barriers CNS 15(1) (2018) 32.

[50] S. Veszelka, A. Toth, F.R. Walter, A.E. Toth, I. Grof, M. Meszaros, A. Bocsik, E. Hellinger, M. Vastag, G. Rakhely, M.A. Deli, Comparison of a Rat Primary Cell-Based Blood-Brain Barrier Model With Epithelial and Brain Endothelial Cell Lines: Gene Expression and Drug Transport, Front Mol Neurosci 11 (2018) 166.

[51] W. Yuan, Y. Lv, M. Zeng, B.M. Fu, Non-invasive measurement of solute permeability in cerebral microvessels of the rat, Microvasc Res 77(2) (2009) 166–73.

[52] L. Shi, M. Zeng, Y. Sun, B.M. Fu, Quantification of blood-brain barrier solute permeability and brain transport by multiphoton microscopy, J Biomech Eng 136(3) (2014) 031005.

[53] J.Y. Li, R.J. Boado, W.M. Pardridge, Blood—Brain Barrier Genomics, Journal of Cerebral Blood Flow & Metabolism 21(1) (2001) 8.

[54] B.J. DeOre, P.P. Partyka, F. Fan, P.A. Galie, CD44 regulates blood-brain barrier integrity in response to fluid shear stress, (2020).

[55] L. Delsing, P. Donnes, J. Sanchez, M. Clausen, D. Voulgaris, A. Falk, A. Herland, G. Brolen, H. Zetterberg, R. Hicks, J. Synnergren, Barrier Properties and Transcriptome Expression in Human iPSC-Derived Models of the Blood-Brain Barrier, Stem Cells 36(12) (2018) 1816–1827.

[56] S. Hamm, B. Dehouck, J. Kraus, K. Wolburg-Buchholz, H. Wolburg, W. Risau, R. Cecchelli, B. Engelhardt, M.P. Dehouck, Astrocyte mediated modulation of blood-brain barrier permeability does not correlate with a loss of tight junction proteins from the cellular contacts, Cell Tissue Res 315(2) (2004) 157–66.

[57] M. Muendoerfer, U.F. Schaefer, P. Koenig, J.S. Walk, P. Loos, S. Balbach, T. Eichinger, C.M. Lehr, Online monitoring of transepithelial electrical resistance (TEER) in an apparatus for combined dissolution and permeation testing, Int J Pharm 392(1-2) (2010) 134–40.

[58] J.D. Siliciano, D.A. Goodenough, Localization of the tight junction protein, ZO-1, is modulated by extracellular calcium and cell-cell contact in Madin-Darby canine kidney epithelial cells, The Journal of Cell Biology 107(6) (1988) 2.

[59] X. Gao, P. Kouklis, N. Xu, R.D. Minshall, R. Sandoval, S.M. Vogel, A.B. Malik, Reversibility of increased microvessel permeability in response to VE-cadherin disassembly, American Journal of Physiology-Lung Cellular and Molecular Physiology 279(6) (2000) L1218–L1225.

[60] L.A. Davidson, B.D. Dzamba, R. Keller, D.W. Desimone, Live imaging of cell protrusive activity, and extracellular matrix assembly and remodeling during morphogenesis in the frog, Xenopus laevis, Dev Dyn 237(10) (2008) 2684–92.

[61] C.J. Sommer, Ischemic stroke: experimental models and reality, Acta Neuropathol 133(2) (2017) 245–261.

[62] G.D. Vatine, R. Barrile, M.J. Workman, S. Sances, B.K. Barriga, M. Rahnama, S. Barthakur, M. Kasendra, C. Lucchesi, J. Kerns, N. Wen, W.R. Spivia, Z. Chen, J. Van Eyk, C.N. Svendsen, Human iPSC-Derived Blood-Brain Barrier Chips Enable Disease Modeling and Personalized Medicine Applications, Cell Stem Cell 24(6) (2019) 995–1005 e6.

[63] Y.B. Arik, M.W. van der Helm, M. Odijk, L.I. Segerink, R. Passier, A. van den Berg, A.D. van der Meer, Barriers-on-chips: Measurement of barrier function of tissues in organs-on-chips, Biomicrofluidics 12(4) (2018) 042218.

[64] F.R. Walter, S. Valkai, A. Kincses, A. Petneházi, T. Czeller, S. Veszelka, P. Ormos, M.A. Deli, A. Dér, A versatile lab-on-a-chip tool for modeling biological barriers, Sensors and Actuators B: Chemical 222 (2016) 1209–1219.

[65] R. Daneman, L. Zhou, A.A. Kebede, B.A. Barres, Pericytes are required for blood–brain barrier integrity during embryogenesis, Nature 468(7323) (2010) 562–566.

[66] D. Bonkowski, V. Katyshev, R.D. Balabanov, A. Borisov, P. Dore-Duffy, The CNS microvascular pericyte: pericyte-astrocyte crosstalk in the regulation of tissue survival, Fluids Barriers CNS 8(1) (2011) 8.

[67] S.I. Ahn, Y. Kim, Human Blood-Brain Barrier on a Chip: Featuring Unique Multicellular Cooperation in Pathophysiology, Trends Biotechnol 39(8) (2021) 749–752.

[68] N. Baeyens, C. Bandyopadhyay, B.G. Coon, S. Yun, M.A. Schwartz, Endothelial fluid shear stress sensing in vascular health and disease, J Clin Invest 126(3) (2016) 821–8.

[69] M. Gerigk, H. Bulstrode, H.H. Shi, F. Tönisen, C. Cerutti, G. Morrison, D. Rowitch, Y.Y.S. Huang, On-chip perivascular niche supporting stemness of patient-derived glioma cells in a serum-free, flowable culture, Lab on a Chip (2021).

[70] J.A. Boos, P.M. Misun, G. Brunoldi, L.A. Furer, L. Aengenheister, M. Modena, N. Rousset, T. Buerki-Thurnherr, A. Hierlemann, Microfluidic Co-Culture Platform to Recapitulate the Maternal-Placental-Embryonic Axis, Adv Biol (Weinh) (2021) e2100609.

[71] C. Lohasz, N. Rousset, K. Renggli, A. Hierlemann, O. Frey, Scalable Microfluidic Platform for Flexible Configuration of and Experiments with Microtissue Multiorgan Models, SLAS Technol 24(1) (2019) 79–95.

[72] J.G. DeStefano, Z.S. Xu, A.J. Williams, N. Yimam, P.C. Searson, Effect of shear stress on iPSC-derived human brain microvascular endothelial cells (dhBMECs), Fluids Barriers CNS 14(1) (2017) 20.

[73] A. Al Ahmad, C.B. Taboada, M. Gassmann, O.O. Ogunshola, Astrocytes and pericytes differentially modulate blood-brain barrier characteristics during development and hypoxic insult, J Cereb Blood Flow Metab 31(2) (2011) 693–705.

[74] Y. Shi, X. Jiang, L. Zhang, H. Pu, X. Hu, W. Zhang, W. Cai, Y. Gao, R.K. Leak, R.F. Keep, M.V. Bennett, J. Chen, Endothelium-targeted overexpression of heat shock protein 27 ameliorates blood-brain barrier disruption after ischemic brain injury, Proc Natl Acad Sci U S A 114(7) (2017) E1243–E1252.

[75] Y. Liu, S.R. Cox, T. Morita, S. Kourembanas, Hypoxia Regulates Vascular Endothelial Growth Factor Gene Expression in Endothelial Cells, Circulation Research 77(3) (1995) 638–643.

[76] B.L. Krock, N. Skuli, M.C. Simon, Hypoxia-induced angiogenesis: good and evil, Genes Cancer 2(12) (2011) 1117–33.

