## Supplementary information for "3D *in vitro* blood-brain-barrier model for investigating barrier insults"

### Supplementary Methods

**Platform fabrication**: The multilayer microfluidic device was designed by using Inventor software (Autodesk, München, Germany). The device featured: one poly(dimethylsiloxane) (PDMS, Sylgard 184, Dow Corning Corp., Midland, MI, USA) microchannel layer that was plasma bonded ( PDC-002, Harrick Plasma, New York, USA) to a 1.5# glass coverslip, which had been patterned with indium-tin-oxide (ITO) electrodes (Glass-ITO, 30-60 Ohms, Diamond Coatings, West Midlands UK); a semipermeable polyethylene terephthalate (PET) membrane with a pore size of 0.4 µm and a thickness of 12 µm (it4ip, Louvain-la-Neuve, Belgium); one PDMS interlayer; a platinum-patterned PET (Sigma-Aldrich, Buchs, Switzerland) electrode layer; a PDMS hanging-drop and medium-reservoir layer.

*Fabrication of PDMS layers:* To form the microchannel layer, the mixed and degassed PDMS (elastomer: curing agent in a 10:1 (w/w) ratio) was spin-coated on a glass slide, which had been previously coated with a sacrificial layer of 5% polyvinyl alcohol (Sigma-Aldrich, Buchs, Switzerland). The PDMS was spin-coated at 650 rpm for 45 s to obtain a 100-µm-thick film and subsequently cured in an oven at 80 °C for 2 h. The cured PDMS film was then cut by using a $CO_2$ laser cutter (Universal Laser Systems, Vienna, Austria) to pattern the geometry of the microchannel. As the PVA is water soluble, the PDMS film was peeled off from the glass slide after immersing the slide in deionized (DI) water to release the microchannel layer, and then dried in an oven at 60 °C for 10 min.

The PDMS interlayer was fabricated by pouring 10 g of mixed and degassed PDMS (elastomer: curing agent in a 10:1 (w/w) ratio) into 145 mm Petri dishes (Greiner Bio-One, Gallen, Switzerland). After curing for 2 h at 80 °C, the PDMS was removed from the dishes, and the brain compartment and inlet /outlet structures were marked with the laser cutter before they were manually cut using a biopsy puncher.

The hanging-drop layer and the open reservoirs were cast from a 3D-printed master mold (Protolabs, Feldkirchen, Germany) by using soft lithography. The mixed and degassed PDMS (elastomer: curing agent in a 10:1 (w/w) ratio) was poured onto the master mold. After curing for 2 h at 80 °C, the PDMS hanging-drop layer and the PDMS open reservoirs were peeled off the master mold and cut into individual chips. The hanging-drop structures were punched through with a 3 mm biopsy punch.

*Electrode fabrication:* The glass-ITO coverslip electrodes were fabricated by using photolithography and wet etching. Positive photoresist S1813 was spin-coated on an ITO-coated coverslip and patterned by using a mask aligner and UV exposure. After resist development, the patterned glass-ITO coverslip was immersed into the etching solution (HCl (36%): $HNO_3$ (67%): $H_2O$ = 10: 1: 36 (v/v/v)) for 120 s at room temperature, followed by photoresist stripping via acetone. The glass-ITO coverslips with patterned ITO electrodes were rinsed in DI water and dried. To reduce the contact resistance between the ITO electrodes and the spring connectors, 270-nm-thick platinum was deposited on the ITO contact pads on the glass-ITO coverslip by using an ion-beam evaporator and patterned with a shadow mask.

The top 270-nm-thick platinum electrodes were deposited on a 6", 150-um-thick PET film by using an ion-beam evaporator and patterned with a shadow mask (Supplementary Fig. 3). After Pt deposition, the PET film was diced into individual PET-Pt electrode slides (15 mm × 21 mm) by using the laser cutter. Three 2 mm-diameter holes were cut by the laser cutter on the electrode slides to realize the openings for the microchannel reservoirs and for the hydrogel compartment.

*Chip assembly:* To bond the glass-ITO electrode layer and the microchannel layer, the glass-ITO substrates were exposed to oxygen plasma (2 min, 30 W using a PDC-002, Harrick Plasma, New York, USA), which was followed by APTES (Sigma-Aldrich) treatment in a vacuum chamber. After being rinsed with DI water, the APTES-treated glass-ITO substrates were bonded with the plasma-treated microchannel layer.

The PET porous membrane was sandwiched between the microchannel layer and the PDMS interlayer by using uncured PDMS as a glue (elastomer: curing agent in a 10:3 (w/w) ratio). The PDMS glue was spin-coated on a glass slide (50 mm × 75 mm) at 3000 rpm for 90 s. The plasma-bonded, glass-ITO-microchannel assembly and the PDMS interlayer were carefully placed on the spin-coated PDMS glue layer to transfer a thin layer of uncured PDMS on the layers to be bonded. Then, the PET membrane and the two PDMS-coated layers were aligned, degassed for 30 min in a vacuum chamber, and cured at 80 °C for 1 h.

The PET-Pt electrode layer was treated with oxygen plasma and then immersed into a 3% APTES solution at 80 °C for 30 min to modify the surface properties. The layer was then bonded between the lower assembly and the hanging-drop layer by using oxygen plasma. Finally, the medium reservoirs were plasma bonded to complete the chip assembly.

*TEER Platform Assembly:* Eight chips were placed between a custom-made printed circuit board (PCB) and a chip holder. The PCB was designed in Altium Designer 17.0 and ordered from PCBWay (Hangzhou, China). Electrical connections between the PCB and the microfluidic devices were obtained by contacting the electrode pads from above by using spring-loaded pins (0956-0-15-20-75-14-11-0, Mill-Max Mfg. Corp., Oyster Bay, USA). The

PCB featured eight rectangular openings (16 mm × 26 mm) to allow for medium refreshing, permeability measurements and visual access to the chips without disassembling the platform.

The microfluidic device holder was custom-made in PMMA, which had been structured by laser cutting. The holder was placed in a well plate (Nunc™ OmniTray™, Thermofisher Scientific, Reinach, Switzerland) for simple handling and compatibility with lab automation equipment. The microfluidic device holder and the well plate featured eight openings that were aligned with the PCB openings for visual examination of the cells by using an inverted confocal microscope. The openings allowed the water immersion objective to contact the bottom coverslip for high-resolution imaging. Finally, the platform was covered with a laser-cut polystyrene lid to prevent medium evaporation from the microfluidic devices and to allow for cable connections to the TEER electrodes.

**Cell culture:** Human cerebral microvascular endothelial cells (hCMEC/D3, Tebu-Bio GmbH, Offenbach, Germany) were cultured in Endothelial Growth Medium (EGM™-2 , Lonza, Basel, Switzerland) in a cell-culture flask, which had been previously coated with 150 μg/mL collagen type I (collagen I, Roche, Basel, Switzerland). All experiments were performed with hCMEC/D3 (ECs) between passage numbers 36-39. Human astrocytes (HAs, ScienCell, Carlsbad, USA) and human pericytes (HPs, ScienCell, Carlsbad, USA) were cultured in 20 μg/mL Poly-l-Lysine (PLL, Sigma-Aldrich, Buchs, Switzerland) coated flasks and maintained in human astrocyte medium (HAM, ScienCell, Carlsbad, USA) and human pericyte medium (HPM, ScienCell, Carlsbad, USA), respectively. The HAs and HPs were used at passage numbers 3-6. All cells were maintained at 37 °C and 5% $CO_2$ inside a cell-culture incubator. The medium was replaced every 2 days. HAs and HPs were detached using 0.05% trypsin/EDTA (Gibco, Thermofisher Scientific, Reinach, Switzerland) and hCMEC/D3 were detached using 0.25% trypsin/EDTA (Gibco, Thermofisher Scientific, Reinach, Switzerland).

For barrier formation, EGM-2 without vascular endothelial growth factor (VEGF) was used, since VEGF has been shown to increase the permeability of endothelial cell monolayers.

Cell metabolic activities under different medium conditions were quantified using a (3-[4, 5-dimethylthiazol-2-yl]-2, 5-diphenyltetrazolium bromide) (MTT reagent, Invitrogen, Thermofisher Scientific, Reinach, Switzerland) assay. ECs, HAs, and HPs were seeded in VEGF-free EGM-2, HAM, and HPM, respectively, in a collagen I- or PLL-coated 96-well plate at a density of $1 \times 10^4$ cells/well. After 48 h of culturing under the respective medium and coating conditions, the medium of six wells of each cell type was changed to the EAPM mixture (VEGF-free EGM-2: HAM: HPM = 1: 1: 1). After 48 h of culturing, 20 μL of MTT reagent

(5mg/mL) were added to each well and incubated in a well plate for 4 h in the cell culture incubator. 100 μL of the solubilization solution were then added to each well, and the optical absorbance at 575 nm of each sample was measured using a Tecan plate reader (Infinite M1000 PRO, Tecan Trading AG, Männedorf, Switzerland).

**3D Cell Culture in the device:** At day -3, the microfluidic devices were sterilized by exposure to ultraviolet (UV) light for 30 min and mounted into the holder and assembled with the PCB. The microchannel and the brain compartment of each chip were coated with 500 μg/mL PLL for 2 h in a cell-culture incubator to improve the adhesion of the hydrogel in the compartments and to promote cell attachment on the porous membrane. The PLL solution was then removed, and the channels and compartments were rinsed with and stored in DI water.

To form a 3D astrocyte-pericyte network, DI water was removed from the chips, and 15 μL of human pericyte suspension at a concentration of $2 \times 10^4$ cells/mL in HPM were loaded into the brain compartment of each chip. The microchannel was also filled with the HPM from the side reservoirs. The HP-loaded devices were placed into a cell culture incubator for 2 h in a static configuration to allow for the HPs to sediment and to attach to the porous membrane. Then, the non-adherent cells were removed from the chips by gently replacing the medium in the brain compartment in the hanging-drop configuration.

The next day, the medium on the chips was removed, and 10 μL of the hydrogel-HA mixture was loaded into the brain compartment of each chip. The hydrogel was prepared by mixing Gamma 2-RGD (Manchester BIOGEL, Cheshire, UK) and collagen I (Advanced Biomatrix, Carlsbad, USA). Prior to mixing, the Gamma 2-RGD was diluted 2.5 times in Peptisol (Manchester BIOGEL) to reduce the hydrogel stiffness. The diluted Gamma 2-RGD was then mixed with collagen I solution to reach a final collagen concentration of 1.6 mg/ml. HAs were then resuspended in the liquid hydrogel mixture at a concentration of $1 \times 10^4$ cells/mL. After loading the hydrogel-HA mixture, the microchannels were filled with EAPM, and the chips were placed in the cell-culture incubator for 10 min in a static configuration to allow for the solidification of the hydrogel. Then, the medium in the microchannel was refreshed with pre-warmed EAPM, and 20 μL of pre-warmed EAPM were loaded into the brain compartments on the hydrogel. The chips were then placed in the incubator, and the medium was exchanged 3 times within the 1$^{st}$ hour to balance the ion concentration in the hydrogel, as per the manufacturer instructions. The cells were afterwards cultured for 2 days on chip to form a 3D astrocyte-pericyte network.

At day 0, the medium in the microchannels was removed, and the channels were washed 3 times with pre-warmed phosphate-buffered saline (PBS). The microchannels were then filled with a 200 μg/mL collagen I solution for 30 min to promote EC attachment in the microchannels and on the porous membranes. The microchannels were rinsed with pre-warmed DI water twice. Subsequently, 80 μL of hCMEC/d3-cell suspension at a concentration of $1 \times 10^7$ cells/mL in EAPM were loaded into the microchannels of each device. The devices were immediately turned upside down to achieve a hanging-drop configuration and incubated for 2 h in the cell-culture incubator to allow for cell sedimentation and attachment to the PET membrane. After 2 h, the non-adherent cells were removed by rinsing the microchannels with fresh EAPM. Afterwards, the platform was placed in a standing-drop configuration on a programmable tilting stage (InSphero AG, Schlieren, Switzerland) in the incubator. The tilting angle was increased gradually every day to allow for barrier formation (Supplementary Fig. 6). Medium exchange was performed daily.

**TEER Measurements:** TEER measurements were performed using an HF2-LI lock-in amplifier (Zürich Instruments AG, Zürich, Switzerland). The microfluidic devices were contacted via the custom-made PCB to route the connections from the lock-in amplifier to the integrated electrodes. A custom-made LabVIEW (National Instruments, Austin, US) user interface was used to control the selection of the microfluidic device and to control signal acquisition. Signal routing on the PCB was performed by the on-board ADG1407 multiplexers (Analog Devices, USA).

A sinusoidal AC voltage signal, with a central frequency that was swept from 500 Hz to 20-kHz and with an amplitude of 200 $mV_p$ (Supplementary Fig. 8), was applied to the working electrode (WE) of the selected microfluidic device, while the CE was kept at pseudo-ground by the HF2-LI transimpedance amplifier (HF2TA, Zürich Instruments AG). The current flowing through the CE, i.e. across the BBB model, was converted to voltage using the HF2TA with a 1-kΩ feedback resistor and sampled by the HF2-LI. Differential voltage measurements between the vascular and brain compartments were acquired via the sensing electrode (SE) and the reference electrode (RE), in close vicinity to the current-injection WE and CE, respectively. Voltage and current measurements were then used to calculate the resistance of the barrier layer.

The initial resistance values ($R_0$) were measured on day 0 before the ECs were seeded and represented the background resistance the system. All the resistance data ($R_{TEER}$) were calculated by subtracting the baseline resistance value ($R_0$) from the measured resistance data

($R_m$) and then normalized with respect to the permeable area of the microchannel and, finally, converted to TEER values ($\Omega \cdot cm^2$) using the following equation (Supplementary Fig. 8):

$$TEER = R_{TEER} \times A = (R_m - R_0) \times A$$

where A indicates the permeable area of the microchannel in $cm^2$.

The alignment of the electrodes with the fluidic network was performed manually, which resulted in differences in baseline resistances and TEER sensitivity among different chips. Therefore, TEER values were normalized between 0% (baseline TEER at day 0, before loading of the ECs) and 100% (TEER value at day 4) to compare different chips and to evaluate the formation and the disruption of the barriers.

**Permeability Measurement:** 4 kDa FITC-dextran (Sigma-Aldrich) was used as a fluorescent tracer to assess the permeability of the BBB models. After 4 days of EC culture in the devices, 200 µL of 200 µg/mL of FITC-dextran in EAPM were loaded into the microchannel, while fresh EAPM was loaded into the hanging-drop compartment. After 5 h of incubation, either in static or dynamic mode according to the BBB model under investigation, 7 µL of medium aliquots were sampled from each microchannel and from the brain compartments. The aliquots were then pipetted in a 384 well plate (Flat Bottom Black Polystyrene, Greiner Bio-One, Gallen, Switzerland) prefilled with 7 µL EAPM. The fluorescence intensity of each sample aliquot at 520 nm was measured with a Tecan microplate reader (490 nm excitation wavelength, 520 nm emission wavelength). A standard curve with different concentrations of FITC-dextran in EAPM was also acquired on the same measurement plate to quantify the concentration of fluorescence tracer in each sample well. Control measurements using microfluidic chips without cells were also acquired for each experimental set.

The apparent permeability coefficient ($P_{app}$) was calculated by using the following equation:

$$P_{app} = \frac{V_{HD} \times C_{HD}}{A \times t \times (C_{MC} \times V_{MC} + C_{HD} \times V_{HD}) / (V_{MC} + V_{HD})}$$

where $P_{app}$ is the permeability coefficient, $t$ is the time of incubation in seconds, $V_{HD}$ is defined by the volume of medium in the hanging-drop compartment, $C_{HD}$ is the measured concentration of FITC-dextran in the hanging-drop compartment at incubation time $t$, $V_{MC}$ is the volume of the microchannel, $C_{MC}$ is the measured concentration of FITC-dextran in the microchannel at time $t$, and $A$ is the permeable area of the microchannel in $cm^2$.

**EDTA treatment and live imaging***:* Ethylenediaminetetraacetate (EDTA; Thermofisher Scientific) was used to disrupt the BBB after 4 days of EC culture in the devices to validate the on-chip TEER sensor and test the live-imaging capabilities of our device in detecting rapid variations in EC-layer organization. Before exposing the cells to EDTA, the culture medium in the microfluidic devices was changed to EAPM with 1 µM SiR-actin (Spirochrome AG, Schaffhausen, Switzerland) and 2 drops/mL Hoechst (Hoechst 33342 NucBlue staining solution, Invitrogen, Thermofisher Scientific) to fluorescently stain the cells for live imaging. After 1 h of incubation, the platform was transferred to a confocal microscope to observe the behavior of the ECs *in situ* by simultaneous live, confocal imaging and real-time TEER detection. Subsequently, the medium in then microfluidic chips was replaced by a pre-warmed 8 mM EDTA-PBS solution. Imaging and TEER-measurement intervals were set to 2 min.

Live imaging was carried out using a Nikon spinning-disk confocal microscope (W1-Sora, Nikon, Egg, Switzerland) with environmental control for live-cell imaging using a 40× water-immersion objective. Image processing was done using the NIS-Elements software package (Nikon, Egg, Switzerland).

**Oxygen/glucose deprivation (OGD) treatment and live imaging***:* After 4 days of EC culturing in the devices and prior to performing OGD studies, the culture medium in the microfluidic devices was changed to EAPM with 5 µM of Image-iT Green Hypoxia Reagent (Thermofisher Scientific), 1 µM SiR-actin and 2 drops/mL Hoechst to stain the cells for live-imaging. After 1 h of incubation, OGD conditions were applied by changing the medium in the chip to Dulbecco's Modified Eagle Medium, with no glucose (DMEM (glucose-free), Gibco, Thermofisher Scientific) and by reducing the $O_2$ level to 0% by placing the platform into an environmental chamber on the confocal microscope stage, which was flushed with 5% $CO_2$, 95% $N_2$ and maintained at 37 $^{\circ}$C. Live imaging was performed using a Nikon spinning disk confocal microscope with a 40× water-immersion objective. Imaging and TEER-measurement sampling intervals were set to 2 min. Image processing was done using the NIS-Elements software package.

**Immunofluorescence and confocal imaging***:* For sample fixation, the cells in the devices were washed with PBS and fixed in intracellular (IC) fixation buffer (Sigma-Aldrich) for 20 min at room temperature (RT), followed by 20 min of 0.1% Triton X-100 (Sigma-Aldrich). After washing and blocking with 5% (w/v) bovine serum albumin (BSA, Sigma-Aldrich) for 2 hours at RT, conjugated antibodies against the tight-junction protein ZO1-1A12 (conjugated with Alexa Fluor 488, 1: 200 dilution, Thermofisher Scientific), the intercellular adherent protein VE-cadherin (VE-CAD, conjugated with Alexa Fluor 647, 1: 50 dilution, BD Biosciences, Allschwil, Switzerland), glial fibrillary acidic proteins (GFAP, conjugated with Cy3, 1:200 dilution, Merck Millipore, Schaffhausen, Switzerland), and alpha smooth muscle actin (α-SMA, conjugated with Cy3, 1:400 dilution, Merck Millipore) were added to a 0.1% (w/v) BSA solution in DPBS and incubated at 4 °C overnight. After washing the samples with PBS, the cells were incubated with the nuclear stain Hoechst for 30 min at RT and stored in PBS at 4 °C until image acquisition. Immunofluorescence images of the barrier were acquired with a Nikon spinning disk confocal microscope using a 40× water-immersion objective. Image processing was done using the NIS-Elements software package.

**Real-time quantitative PCR (RT-qPCR):** To compare the relative gene expression levels among different treatment and culturing groups, samples for RT-qPCR analysis were generated by harvesting ECs cultivated on chips in mono- or co-culture, static or under shear stress, and with or without hypoxia treatment (n=3 technical replicates and n=3 biological replicates). RNA isolation was performed with the NucleoSpin RNA XS kit according to the manufacturer's instructions (Machery-Nagel, Oensingen, Switzerland). For complementary deoxyribonucleic acid (cDNA) synthesis, the isolated RNA was reversely transcribed into a 20 µL total volume using the RNA to cDNA EcoDry™ Premix (Oligo dT) (TaKaRa, Saint-Germain-en-Laye, France) according to the manufacturer's instructions. RT-qPCR was conducted using a QuantStudio 3 (Applied Biosystems, Thermofisher Scientific, Reinach, Switzerland), and primers were purchased from Integrated DNA Technologies (IDT, Zürich, Switzerland). The gene expression assays with human-specific primers for tight junction protein ZO-1 (TJP1), adherent junction protein VE-cadherin (CDH5), vascular endothelial growth factor A (VEGFa), TFRC (CD71) were used for gene expression analysis in human ECs (TableS 1). X-fold changes in the relative mRNA expression were determined using the comparative cycle method ($2^{-\Delta\Delta Ct}$) normalized to the housekeeping gene TFRC, since the TFRC genes were stable under different conditions[1, 2].

**Numerical Simulations:** The layout of the ITO electrode was modeled and optimized by using COMSOL (COMSOL Multiphysics 5.3a, COMSOL Multiphysics GmbH, Zürich, Switzerland).

*Fluid dynamics:* To estimate the fluid dynamics through the microfluidic devices, the CFD module of COMSOL was used. The fluid flow through the microfluidic device was set as a laminar flow, since the microfluidic channel featured a low Reynold's number.[3] The properties of water (at 37°C) were used to estimate those of the media in the experimental setup. The fluid was considered incompressible, and the dynamic viscosity µ was set to 0.7 mPa·s.[4] From simulations, we obtained a hydraulic channel resistance of

$$R_{hydro} = 5.53 \times 10^{10}\ kg/(m^4 \cdot s)$$

After calculating the channel resistance, we used this parameter to estimate the pressure drop across the channel during tilting and to calculate the resulting flow rate in MATLAB. The inlet flow rate in the COMSOL simulation was then set to the calculated flow rate, as shown in Supplementary Figure 6, to estimate the shear stress on the endothelial cell layer.

*Current density:* To estimate the current density across the cellular barrier, the secondary Current Distribution (CD) module was used in COMSOL. The properties of water (at 37°C) were used to estimate those of the media in the experimental setup with an electrolyte conductivity of 1.6 S/m[5]. The electrical double layer capacitances of the WE and SE were set to 0.0337 F/m$^2$, and those of the CE and RE were set to 0.25 F/m$^2$.[6, 7] Frequency domain perturbation studies were used to simulate the current density at frequencies from 10 Hz to 2000 kHz. Additional simulation results of the current distribution in the barrier area are shown in Supplementary Fig. 3.

*Fluid-flow modeling:* The flow rate through the microchannel and the shear stress on the permeable barrier area were determined numerically using MATLAB. The custom-made MATLAB script considered the hydraulic resistance of the microchannel, which was estimated in COMSOL, and the shear stress across the permeable area of the porous membrane Supplementary Fig. 2, 6). The shear stress distribution in the barrier area was simulated by using COMSOL, using the flow and pressure values calculated in MATLAB as input parameters.

The cell culture medium is incompressible, and the medium flow in the microchannel is considered a laminar flow. The laminar flow is driven by the height difference ($\Delta h$) of the medium levels in the two side reservoirs, which generates hydrostatic pressure differences ($\Delta P$) in the microchannel. The hydraulic resistance ($R_{hydro}$) of the microchannel has been calculated. The volumetric flow rate ($Q$) can be expressed as:

$$Q = \frac{\Delta P}{R_{hydro}} \quad (1)$$

The microchannel hydraulic resistance ($R_{hydro}$) was calculated by using COMSOL Multiphysics:

$$R_{hydro} = 5.53 \times 10^{10}\ kg/(m^4 \cdot s) \quad (2)$$

The hydrostatic pressure difference ($\Delta P$) is controlled by the height difference ($\Delta h$) of the medium levels in the reservoirs:

$$\Delta P = \rho g \Delta h \quad (3)$$

As shown in Fig. S6, $\Delta h$ is determined by the height difference ($\Delta h_r$) of the two reservoirs upon tilting and the difference in the medium height in the reservoirs ($h_1 - h_2$):

$$\Delta h = \Delta h_r + (h_1 - h_2) \tag{4}$$

$\Delta h_r$ is expressed as:

$$\Delta h = l \cdot \sin\theta \tag{5}$$

$h_1 - h_2$ is expressed as:

$$h_1 - h_2 = -\frac{2 \cdot \int_0^t Q\,dt}{A} \tag{6}$$

Here, A is the cross-sectional area of the medium reservoir.

Therefore, the volumetric flow rate ($Q$) can be expressed by:

$$Q = \frac{\rho g \left(L \sin\theta - \frac{2\int_0^t Q dt}{A}\right)}{R_{hydro}} \tag{7}$$

The flow rate ($Q$) was calculated numerically using MATLAB.

**Statistical Analysis***:* All results are presented as mean values ± standard deviation (s.d.). Statistical analysis was performed using Prism GraphPad 9. Two-way ANOVA with Tukey's multiple comparisons test was used for two-groups comparisons along with a t-test or one-way ANOVA (**p* < 0.05, ***p* < 0.01, and ****p* < 0.001, *****p* < 0.0001).

### Supplementary Figures

### BBB platform

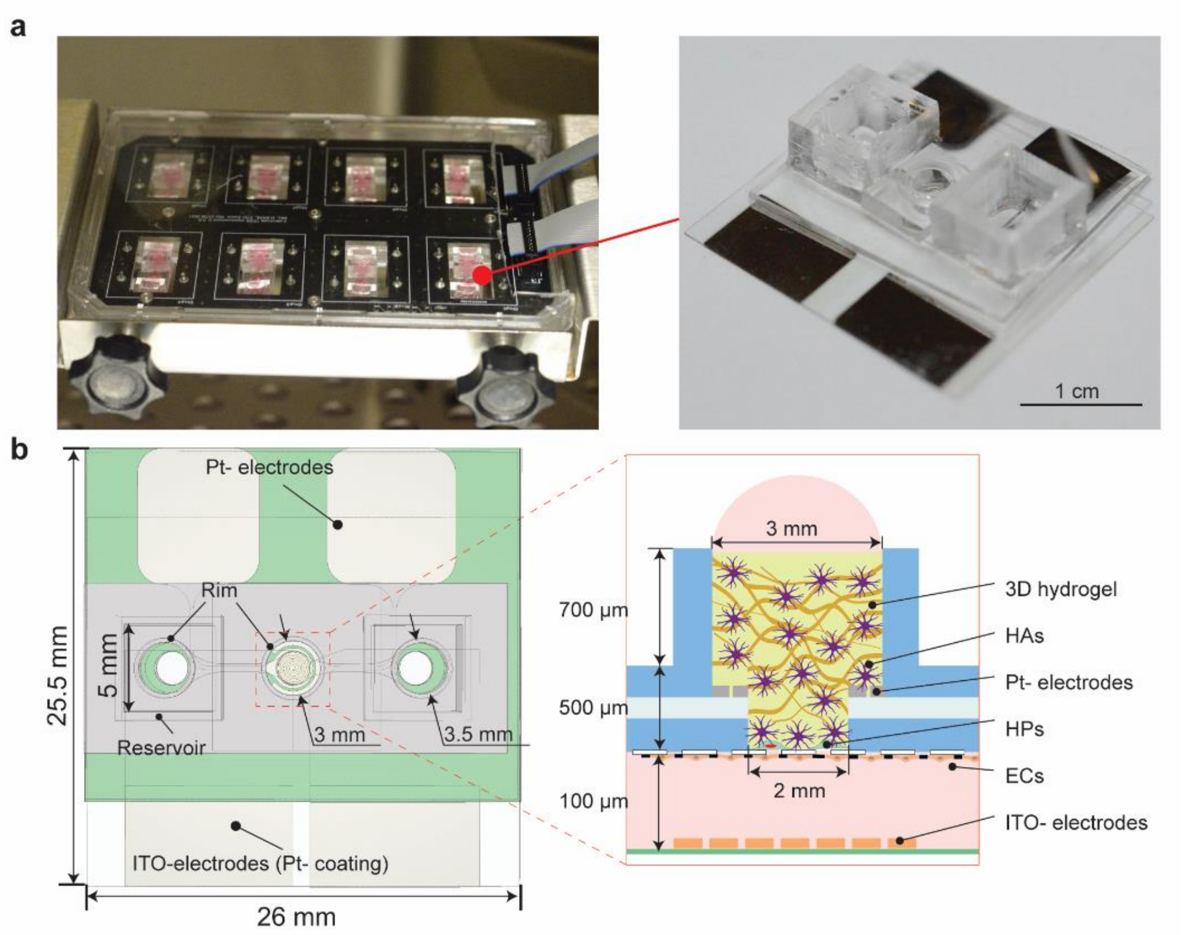


Figure S1. Design and dimensions of the BBB chip. a) Photograph of the BBB platform with 8 chips (left) and of a single chip (right). The BBB platform was composed of a top PCB to contact the on-chip electrode pads and a bottom layer to maintain the chips in position. Each chip featured four, Pt-coated electrode pads and large medium reservoirs; b) Design and dimensions of the BBB microfluidic chip (top view) and cell organization in the chip (side view). Cell-culture medium (in pink) was flown through the microfluidic channel and loaded on top of the brain compartment, which was filled with 3D hydrogel (yellow).

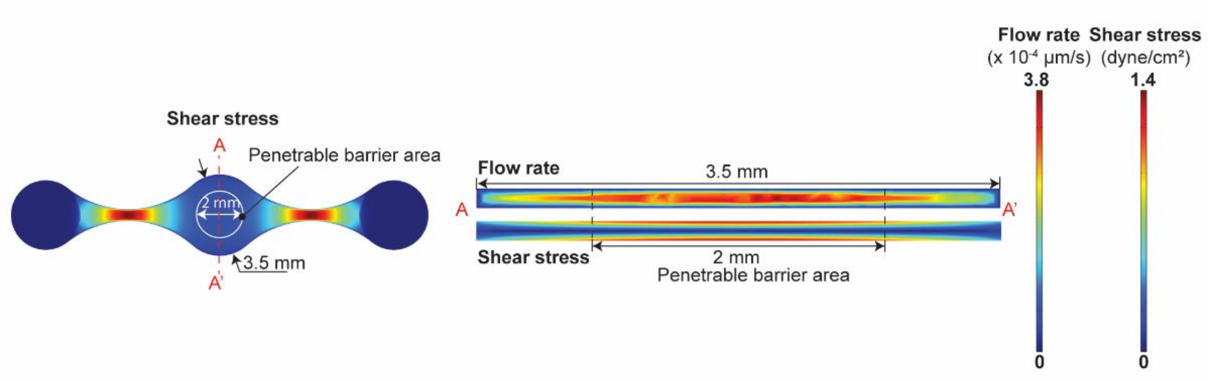


Figure S2. Simulation of shear stress using the COMSOL Multiphysics software. The diameter of the barrier area, which connected the vascular and brain compartments, was designed smaller than the total channel width so as to achieve uniform shear stress on the ECs forming the barrier.

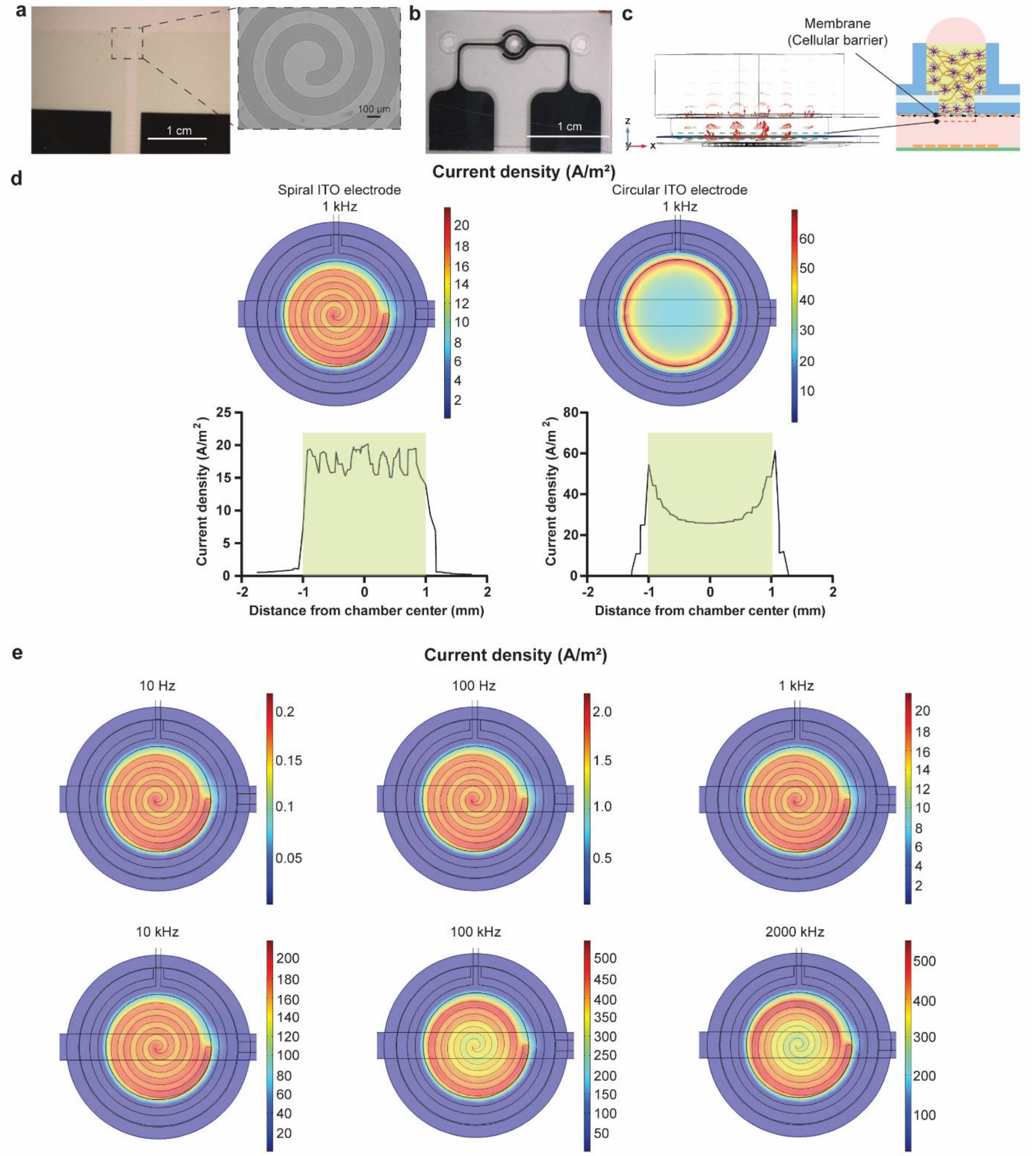


Figure S3. a) Photograph of the glass-ITO electrodes featuring large Pt contact pads. The electrodes were fabricated using transparent ITO on a 1.5# glass coverslip to enable high-resolution imaging. The Pt coating on the contact pads provided reliable electrical connections to external pins and reduced the overall system impedance. The working electrode in ITO featured spiral shape to obtain high uniformity current density across the barrier; b) Photograph of the PET-Pt top ring electrodes; c) FEM simulation of the current density in the chip. A sinusoidal voltage with different carrier frequencies was applied to the ITO working electrode. The red arrows indicate the current distribution from stimulation to sensing electrodes. The current passed through the membrane where the cellular barrier was formed and reported on the tightness of the barrier; d) Comparison of current density at the membrane layer using a spiral or circular working electrode along with the ring-based counter electrode. The spiral working electrode provided higher current uniformity across the cellular barrier at 1 kHz. The simulation was carried out by applying a 200-mV-amplitude sinusoidal stimulation. The plots indicate the root mean squared current density. The porous BBB area connecting the vessel and brain side is indicated through the shaded areas in the graph; e) FEM simulation of the current density across the membrane layer at different frequencies, obtained by applying a sinusoidal voltage stimulus of 200 mV amplitude. For signals below 100 kHz, the current density showed high uniformity across the porous membrane. However, for low frequencies (<1 kHz), the current densities were very low, which rendered signal detection difficult. All TEER values were calculated at 1 kHz. All FEM simulations were carried out using COMSOL Multiphysics.

### Experimental setup

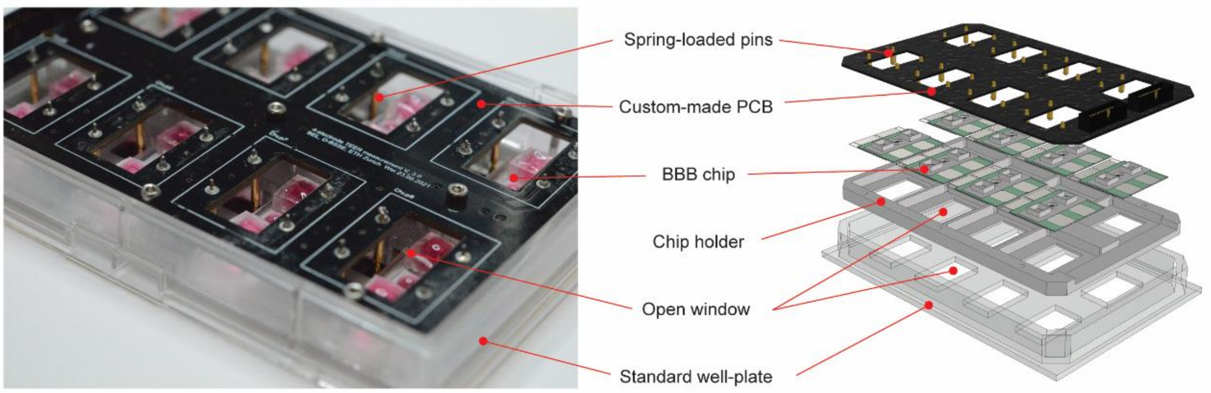


Figure S4. Assembly of the BBB platform. The platform is modular. 8 chips can be mounted between the connecting PCB and the chip holder. The spring-loaded pins on the PCB are aligned and pushed onto the electrode contact pads of the chips upon assembling the platform. The platform was arranged in a standard well-plate format for compatibility with laboratory automation and microscopy tools. Open windows in the PCB were aligned with the microfluidic-chip imaging areas so as to ensure continuous optical access. Open windows in the holder enabled the use of water immersion to allow for use of immersion objectives for high-resolution imaging.

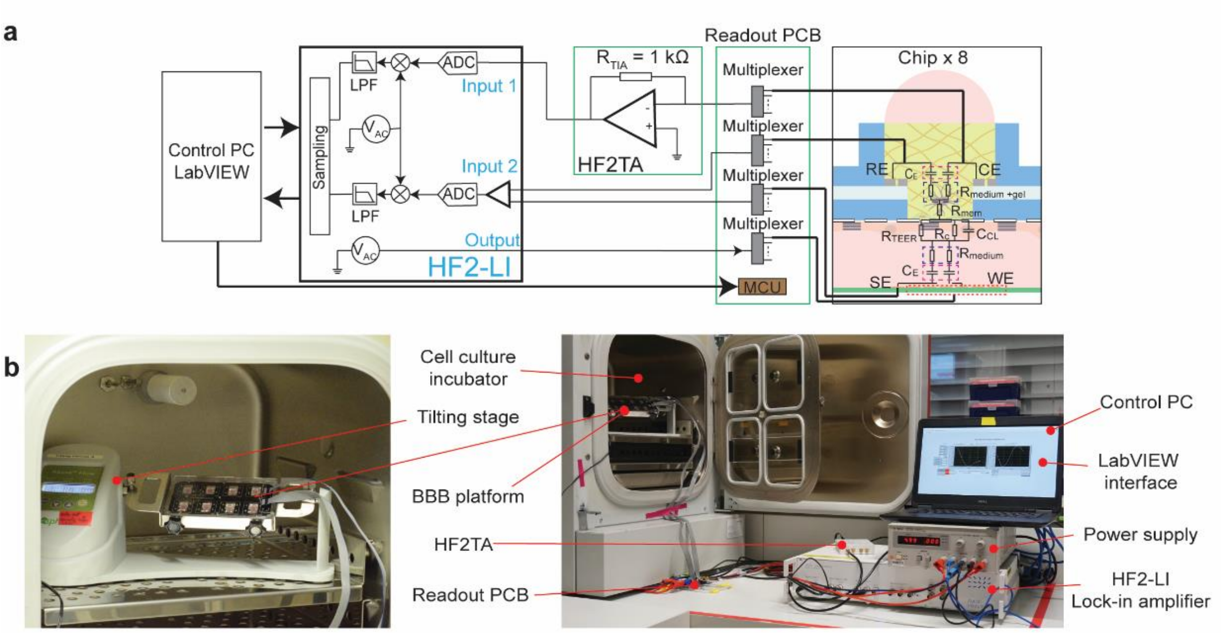


Figure S5. a) The electrical equivalent circuit of the experimental setup. A lock-in amplifier generates the AC stimulation signal, which is then routed to the chips by a custom-made PCB, where a microcontroller (MCU) is used to control the on-board multiplexers. The circuit is then completed by a HF2-LI transimpedance amplifier (HF2TA), which provides current-to-voltage (I-V) conversion. The HF2TA is then connected to the input of the lock-in amplifier to measure the current flowing in the system. The voltage drop across the barrier is measured using a set of dedicated electrodes on chip and by using a differential detection scheme. The input signals in the lock-in amplifier are digitalized (analog-to-digital converter, ADC) , low-pass filtered (LPF), sampled, and subsequently recorded on a control PC; b) Photograph of the experimental setup, showing the BBB platform on the tilting stage in the incubator (left) and the TEER acquisition system (right).

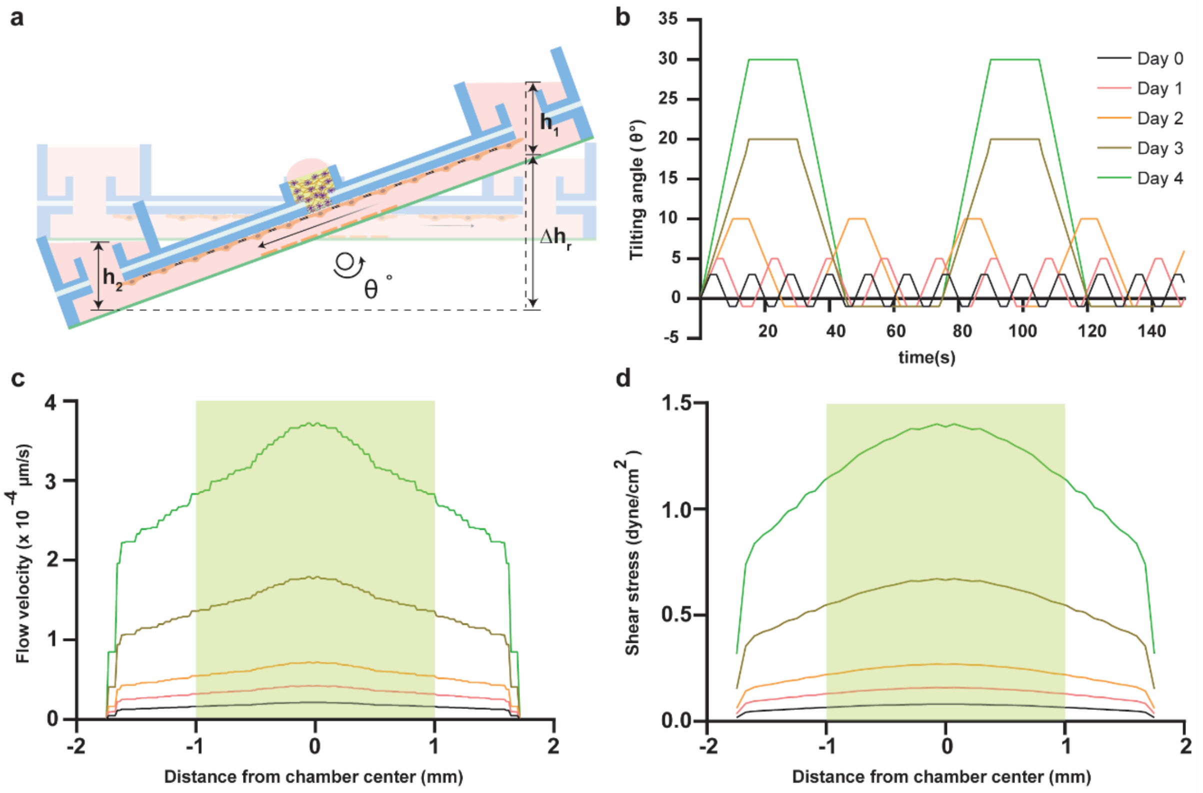


Figure S6. a) Tilting operation of the BBB platform for pump-free flow and dynamic culturing. Medium flow in the eight devices was generated by tilting the platform around an axis perpendicular to the microchannels; b) The tilting profile over time. To avoid damaging the EC layer during barrier formation, the flow rate was gradually increased every day by a stepwise increase of the tilting angles. An asymmetric tilting profile, with large positive tilting angles and small negative tilting angles, was selected to avoid equal-volume, bi-directional flow on the ECs; c) The flow rate at the location of the ECs in the middle of the barrier. The microchannel was wider than the barrier area to avoid having ECs exposed to extremely low flow rates; d) The corresponding shear stress on the ECs. The shaded areas in plots c) and d) indicate the porous surface connecting the vessel and the brain side of the BBB model.

### 3D Blood Brain Barrier

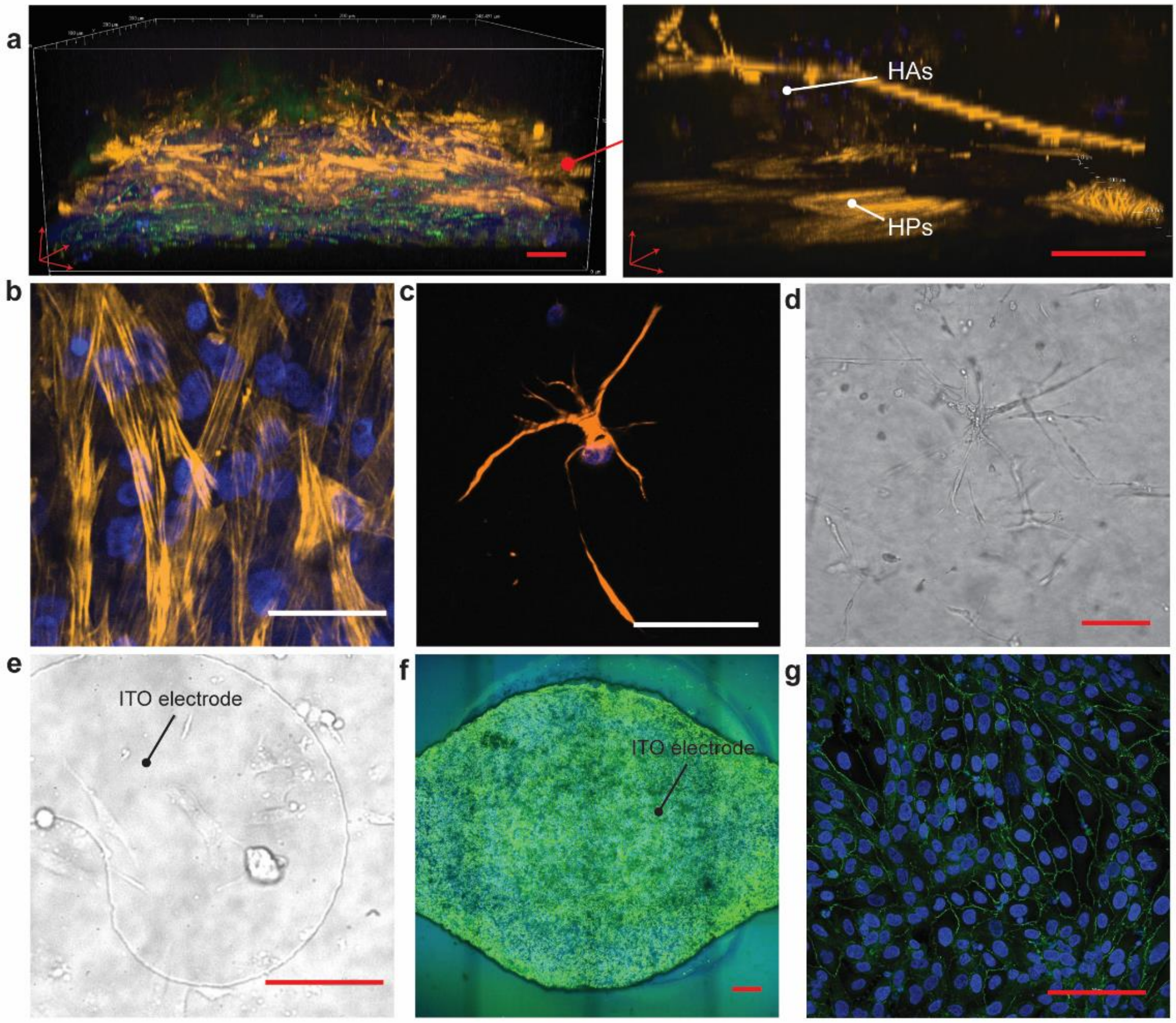


Figure S7. a) The multilayered 3D structure of the on-chip BBB. Endothelial cells formed a continuous monolayer at the bottom side of the porous membrane, and HPs grew on the opposite side of the porous membrane. HAs developed long branches, and the terminals of their processes reached the HPs; b) HPs grew adherent on the porous membrane; c, d) HAs in the hydrogel with GFAP and Hoechst staining (c) and a phase-contrast image (d) showing the characteristic star-like shape of HAs. e) Because of the hanging-drop operation of the chip during EC seeding, few ECs adhered to and grew on the transparent ITO electrodes; f-g) Immunofluorescence staining of the entire cellular barrier for tight junctions (ZO-1) and cell nuclei (Hoechst) in the microfluidic device, which shows that a continuous cell layer was formed across the entire BBB compartment. Scale bars: 100 µm (in red), 50 µm (in white).

### TEER recording on chip

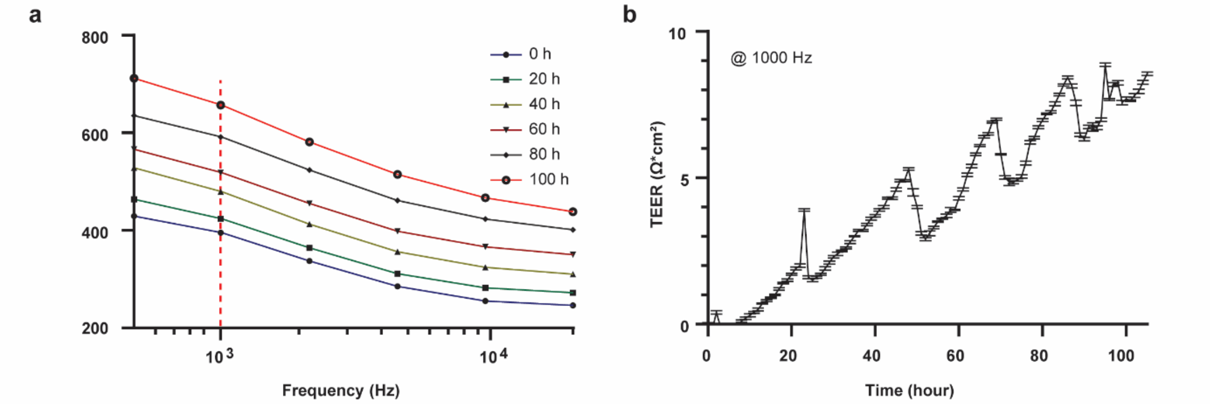


Figure S8. a) Example of impedance spectra, recorded during the BBB formation under dynamic coculture conditions. The impedance spectra were acquired in a range between 500 Hz and 20 kHz. 1 kHz impedance values were selected to calculate the TEER values, as shown in b; b) TEER values were measured with the integrated sensor every hour under dynamic coculture conditions at 1 kHz. Four measurements were acquired per each time point. The graph shows mean values ± s.d..

### Live, confocal imaging of the EC layer on chip under OGD conditions

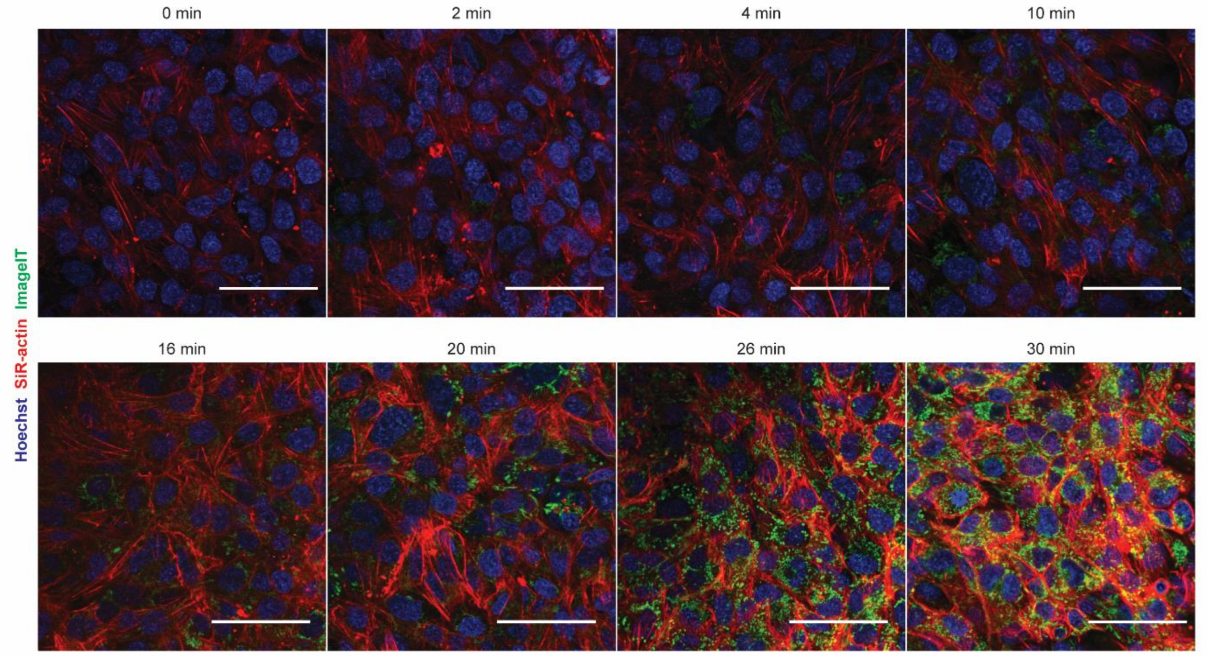


Figure S9. Live fluorescence images of the EC layer during OGD exposure. After 4 minutes of OGD exposure, the fluorescence signal of the hypoxia marker (ImageIT, green) appeared, which indicated that ECs had been under hypoxic stress. After that, the fluorescence signal of the hypoxia marker gradually increased, while F-actin (SiR-actin, red) began to be overexpressed, and actin filaments were rearranged. Strong actin signal, indicating the formation of stress fibers in the ECs, appeared after ~30 minutes of OGD exposure. In blue, cell nuclei (Hoechst). Scale bars: 50 µm.

### Supplementary Tables

Primer sequence for RT-qPCR

Table S1. Primer sequences used for qPCR

| Genes | Forward (5'-3') | Reverse (5'-3') | GenBank ID |
|---|---|---|---|
| ZO-1 | TGCTGAGTCCTTTGGTGATG | AATTTGGATCTCCGGGAAGAC | NM_003257 |
| VE-cadherin | AAACACCTCACTTCCCCATC | ACCTTGCCCACATATTCTCC | NM_001795 |
| VEGFa | AGTCCAACATCACCATGCAG | TTCCCTTTCCTCGAACTGATTT | NM_001025366 |
| TFRC | ACTTGCCCAGATGTTCTCAG | GTATCCCTCTAGCCATTCAGTG | NM_003234 |

### Supplementary Video

Video S1. Live, confocal imaging of the EC layer on chip under OGD conditions

### Supplementary References

[1] S. Bakhashab, S. Lary, F. Ahmed, H.J. Schulten, A. Bashir, F.W. Ahmed, A.L. Al-Malki, H.S. Jamal, M.A. Gari, J.U. Weaver, Reference genes for expression studies in hypoxia and hyperglycemia models in human umbilical vein endothelial cells, G3 (Bethesda) 4(11) (2014) 2159-65.
[2] W. Ju, A.O. Smith, T. Sun, P. Zhao, Y. Jiang, L. Liu, T. Zhang, K. Qi, J. Qiao, K. Xu, L. Zeng, Validation of Housekeeping Genes as Reference for Reverse-Transcription-qPCR Analysis in Busulfan-Injured Microvascular Endothelial Cells, Biomed Res Int 2018 (2018) 4953806.
[3] H.A. Stone, A.D. Stroock, A. Ajdari, Engineering Flows in Small Devices: Microfluidics Toward a Lab-on-a-Chip, Annual Review of Fluid Mechanics 36(1) (2004) 381-411.
[4] A.D. Wong, M. Ye, A.F. Levy, J.D. Rothstein, D.E. Bergles, P.C. Searson, The blood-brain barrier: an engineering perspective, Front Neuroeng 6 (2013) 7.
[5] G. Pucihar, T. Kotnik, M. Kandušer, D. Miklavčič, The influence of medium conductivity on electropermeabilization and survival of cells in vitro, Bioelectrochemistry 54(2) (2001) 8.
[6] J. Martinez, A. Montalibet, E. McAdams, M. Faivre, R. Ferrigno, Effect of electrode material on the sensitivity of interdigitated electrodes used for Electrical Cell-Substrate Impedance Sensing technology, 2017 39th Annual International Conference of the IEEE Engineering in Medicine and Biology Society (EMBC) (2017) 3.
[7] B. Fan, B. Wolfrum, J.T. Robinson, Impedance scaling for gold and platinum microelectrodes, J Neural Eng 18(5) (2021).
